# Quality Control for Single Cell Analysis of High-plex Tissue Profiles using CyLinter

**DOI:** 10.1101/2023.11.01.565120

**Authors:** Gregory J. Baker, Edward Novikov, Ziyuan Zhao, Tuulia Vallius, Janae A. Davis, Jia-Ren Lin, Jeremy L. Muhlich, Elizabeth A. Mittendorf, Sandro Santagata, Jennifer L. Guerriero, Peter K. Sorger

## Abstract

Tumors are complex assemblies of cellular and acellular structures patterned on spatial scales from microns to centimeters. Study of these assemblies has advanced dramatically with the introduction of high-plex spatial profiling. Image-based profiling methods reveal the intensities and spatial distributions of 20-100 proteins at subcellular resolution in 10^3^–10^7^ cells per specimen. Despite extensive work on methods for extracting single-cell data from these images, all tissue images contain artefacts such as folds, debris, antibody aggregates, optical aberrations and image processing errors that arise from imperfections in specimen preparation, data acquisition, image assembly, and feature extraction. We show that these artefacts dramatically impact single-cell data analysis, obscuring meaningful biological interpretation. We describe an interactive quality control software tool, CyLinter, that identifies and removes data associated with imaging artefacts. CyLinter greatly improves single-cell analysis, especially for archival specimens sectioned many years prior to data collection, such as those from clinical trials.

## INTRODUCTION

Tissues are complex assemblies of many cell types whose proportions and properties are controlled by cell-intrinsic molecular programs and interactions with the tumor microenvironment. Recently developed highly multiplexed tissue imaging methods (e.g., MxIF, CyCIF, CODEX, 4i, mIHC, MIBI, IBEX, and IMC)^1–7^ have made it possible to collect single-cell data on 20-100 proteins and other biomolecules in preserved 2D and 3D tissue microenvironments^4,8–11^. Such data are powerful complements to data obtained using dissociative methods such as scRNA-Seq^12–14^. Imaging approaches compatible with formaldehyde-fixed, paraffin-embedded (FFPE) specimens are particularly powerful because they can tap into large archives of human biopsy and resection specimens^15,16^ and also assist in the study of mouse models of disease^17^.

Generating single cell data from high-plex images requires segmenting images^18^ to produce single-cell “spatial feature tables” that are analogous to count tables in scRNA-Seq^18^. In their simplest form, each row in a spatial feature table contains the X,Y coordinate of a cell (commonly the centroid of the nucleus) and integrated signal intensities for each protein marker^19^. Cell types (e.g., cytotoxic T cells immunoreactive to CD45, CD3 and CD8 antibodies) are then inferred from these tables and spatial analysis is performed to identify recurrent short- and long-range interactions significantly associated with an independent variable such as drug response, disease progression, or genetic perturbation.

High-plex spatial analysis has been performed using both tissue microarrays (TMAs), which comprise 0.3 to 1.5 mm diameter “cores” (∼10^4^ cells) from dozens to hundreds of clinical specimens arrayed on a slide, and whole-slide imaging, which can involve areas of tissue as large as 4-6 cm^2^ (∼10^7^ cells). Whole slide imaging is an FDA requirement^20^ for clinical diagnosis, research, and spatial power^21^, but TMAs are nonetheless in widespread use. In this paper, we show that accurate processing of images from both types of specimens is complicated by the presence of imaging artefacts such as tissue folds, slide debris (e.g., lint), and staining artefacts. The problem impacts all data we have examined but is particularly acute with specimens stored for extended periods on glass slides. In our study, this scenario is represented by 25 specimens from the TOPACIO clinical trial of *Niraparib in Combination with Pembrolizumab in Patients with Triple-negative Breast Cancer or Ovarian Cancer* (NCT02657889)^22^, which was completed in 2021. We demonstrate the impact of artefacts on analysis of CyCIF images of TOPACIO tissue specimens and high-plex CyCIF, CODEX, and mIHC datasets from several recently published studies. We then develop a human-in-the loop approach to remove single-cell data affected by microscopy artefacts using a software tool, CyLinter (code and documentation at https://labsyspharm.github.io/cylinter/), that is integrated into the Python-based Napari image viewer^23^. We demonstrate that CyLinter can salvage otherwise uninterpretable multiplex imaging data, including those from the TOPACIO trial. Finally, we demonstrate progress on a deep-learning (DL) model for automated artefact detection; libraries of artefacts identified using CyClinter represent ideal training data for this model. Our findings suggest that artefact removal should be a standard component of processing pipelines for image-based spatial profiling data.

## RESULTS

### Identifying recurrent image artefacts in multiplex IF images

To categorize imperfections and image artefacts commonly encountered in high-plex images of tissue, we examined seven datasets collected using three different imaging methods: (1) 20-plex CyCIF^24^ images of 25 triple-negative breast cancer (TNBC) specimens collected from TOPACIO clinical trial patients^22^; (2) a 22-plex CyCIF image of a colorectal cancer (CRC) resection^21^; (3) a 21-plex CyCIF TMA dataset^25^ comprising 123 healthy and cancerous tissue cores; (4) two 16-plex CODEX^26^ images of a single head and neck squamous cell carcinoma (HNSCC) specimen; (5) a 19-plex mIHC^27^ image of normal human tonsil^25^; (6) 59-plex and (7) 54-plex independent CODEX images of normal large intestine (Supplementary Fig. 1a-g and **Supplementary Table 1**). Raw image tiles were processed using MCMICRO^28^ to generate stitched and registered multi-tile image files and their associated single-cell spatial feature tables. Single-cell data were visualized as UMAP embeddings clustered with HDBSCAN—an algorithm for hierarchical density-based clustering^29^. Images were also inspected by experienced microscopists and board-certified pathologists to identify imaging artefacts.

All specimens comprised 5 µm-thick tissue sections mounted on slides in the standard manner. This involves cutting FFPE blocks with a microtome and floating sections on water prior to capturing them on glass slides. Even in the hands of skilled histologists, this process can introduce folds in the tissue. We identified multiple instances of tissue folds in whole-slide and TMA specimens (**Fig. 1a**, Extended Data Fig. 1a and **Online Supplementary Fig. 1a**). Moreover, we found that cells within tissue folds gave rise to discrete clusters in UMAP feature space due to higher-than-average signals relative to unaffected regions of tissue (**Fig. 1a,b**, Extended Data Fig. 1a,b).

**Fig. 1.**
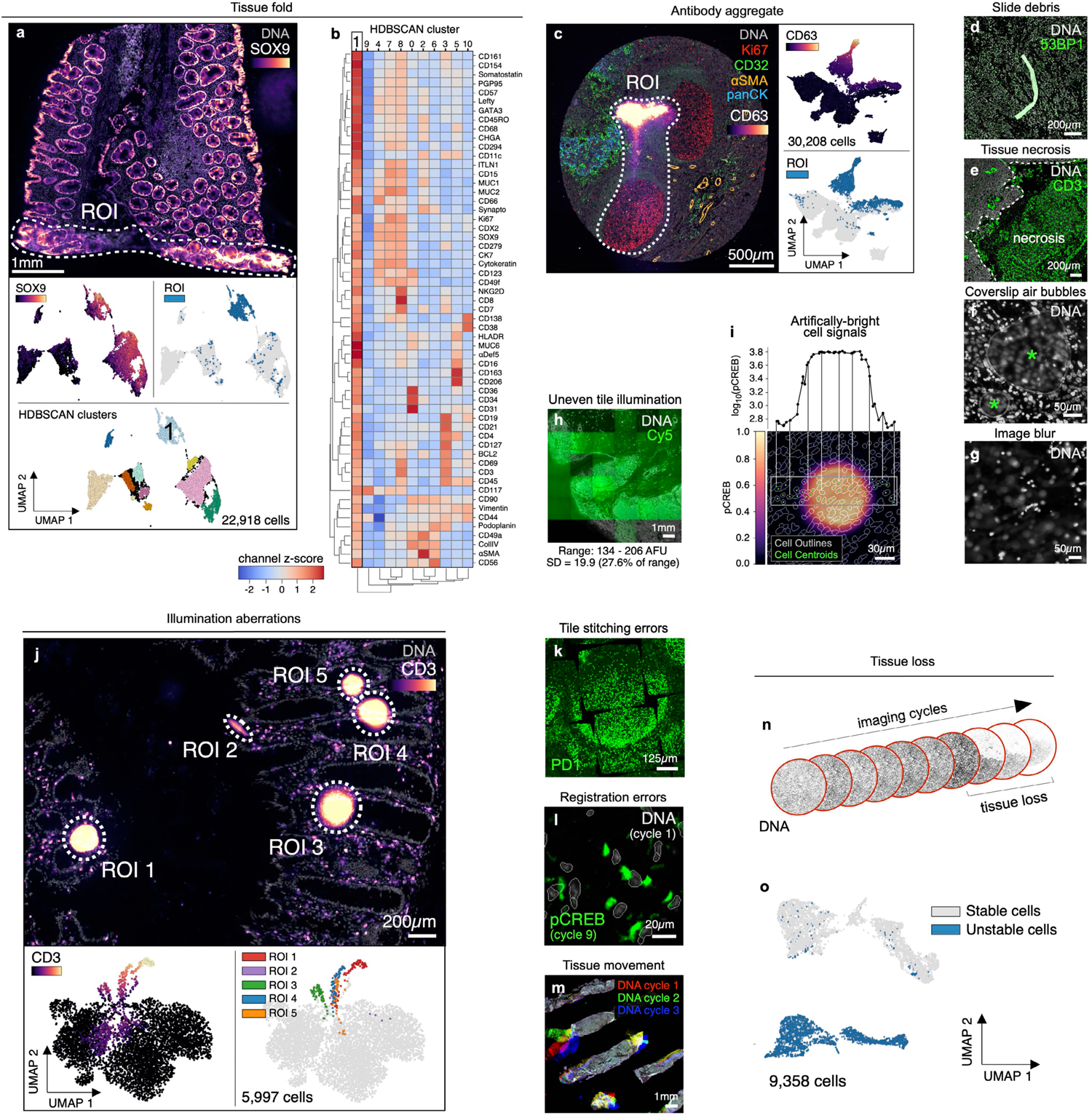
Recurring artefacts in whole slide immunofluorescence images of tissue and their effects on tissue-derived single-cell data. **a**, Top: Field of view from Dataset 6 (large intestine, CODEX, specimen 1) with a tissue fold (ROI, dashed white outline) as viewed in channels SOX9 (colormap) and Hoechst (gray). Bottom: UMAP embedding of 57-channel single-cell data from the image above colored by SOX9 intensity (top left), cells contained fall within the ROI (top right), and HDBSCAN cluster (bottom center). Cluster 1 cells (labeled) are those affected by the tissue fold and form a discrete cluster in UMAP space. **b**, Clustered heatmap showing channel z-scores for HDBSCAN clusters from panel (a) demonstrating that cluster 1 cells (those affected by the tissue fold) are artificially bright for all channels presumably due to a combination of tissue overlap and insufficient antibody washing. **c**, Left: Antibody aggregate in the CD63 channel (colormap) of Dataset 3 (EMIT TMA, core 68, normal tonsil). Hoechst (gray), Ki67 (red), CD32 (green), αSMA (orange), and panCK (blue) are shown for context. Right: UMAP embedding of 20-channel single-cell data from the image shown at left colored by CD63 intensity (top) and whether cells fall within the ROI (bottom). **d**, Autofluorescent fiber in Dataset 1 (TOPACIO, specimen 128) as seen in channels 53BP1 (green) and Hoechst (gray). **e**, Necrosis in a region of tissue from Dataset 1 (TOPACIO, specimen 39) as seen in the CD3 channel (green). **f**, Coverslip air bubbles (green asterisks) in Dataset 1 (TOPACIO, specimen 48) as seen in the Hoechst channel (gray). **g**, Out-of-focus region of tissue in Dataset 1 (TOPACIO, specimen 55) as seen in the Hoechst channel (gray). **h**, Uneven tile illumination in Dataset 4 (HNSCC, CODEX, section 1) as seen in an empty Cy5 channel (green); Hoechst (gray) shown for tissue context. The standard deviation among per-tile median signal intensities was 19.9 arbitrary fluorescence units (AFU), 27.6% of the range (134-206 AFU). **i**, Bottom: Illumination aberration in the pCREB channel (colormap) of Dataset 3 (EMIT TMA, core 95, dedifferentiated liposarcoma) with nuclear segmentation outlines (translucent contours) shown for reference. Top: Line plot demonstrating that artificial pCREB signals of single cells affected by the aberration reach an order of magnitude above background. **j**, Top: Field of view from Dataset 7 (large intestine, CODEX, specimen 2) showing five illumination aberrations (ROIs, dashed white outlines) as viewed in channels CD3 (colormap) and Hoechst (gray). Bottom: UMAP embedding of 52-channel single-cell data from the image above colored by CD3 intensity (left) and whether the cells fall within one of the five different ROIs (right). **k**, Tile stitching errors in Dataset 5 (mIHC, normal human tonsil) as seen in the PD1 (green) channel. **l**, Cross-cycle image registration error in Dataset 3 (EMIT TMA, core 64, leiomyosarcoma) as demonstrated by the superimposition of cycle 1 Hoechst signal (gray) and cycle 9 pCREB signal (green). **m**, Cross-cycle tissue movement in Dataset 1 (TOPACIO, specimen 80) as demonstrated by the superimposition of Hoechst signals from three different imaging cycles: 1 (red), 2 (green), 3 (blue). **n**, Progressive tissue loss in Dataset 3 (EMIT TMA, core 1, normal kidney cortex) across 10 imaging cycles as observed in the Hoechst channel (gray) where overt tissue loss can be seen by cycle 8. **o**, UMAP embedding of cells from Dataset 3 (EMIT TMA, core 1, normal kidney cortex) colored by whether cells remained stable (gray data points) or became detached (blue data points) over the course of imaging demonstrating that unstable cells form discrete clusters in UMAP space.

Bright antibody aggregates were common and also formed discrete clusters in UMAP space (**Fig. 1c**), as were debris in the shape of lint fibers and hair (**Fig. 1d** and **Online Supplementary** Fig. 1b). Despite having relatively low numbers of segmented cells, regions of necrotic tissue also exhibited high levels of background antibody labeling (**Fig. 1e**). Some specimens contained air bubbles likely introduced when coverslips were overlayed on specimens prior to imaging (**Fig. 1f** and **Online Supplementary** Fig. 1c). In principle, artefacts such as tissue folds and air bubbles can be reduced by skilled experimentalists, but access to the original tissue blocks is required.

Additional artefacts were introduced at the time of image acquisition. These included out-of-focus image tiles due to sections not lying completely flat on the slide (**Fig. 1g** and **Online Supplementary** Fig. 1d), fluctuations in background intensity between image tiles (**Fig. 1h**), and miscellaneous aberrations that significantly increased signal intensities over image background (**Fig. 1i**) and generated discrete clusters in UMAP space (**Fig. 1j** and Extended Data Fig. 1c). In some cases, removal of artefacts revealed more subtle problems such as the presence of cells stained non-specifically by all antibodies (e.g., in CODEX Dataset 6; Extended Data Fig. 1d,e). Errors were also observed in tile stitching (**Fig. 1k**) and registration (**Fig. 1l**); in some cases, these problems can be addressed by reprocessing the data, but over-saturation of nuclear stain used for stitching and registration can limit accuracy of even reprocessed data.

Some artefacts were specific to cyclic imaging methods such as CyCIF^24,30^, CODEX^26^, and mIHC^27^ that generate high-plex images through multiple rounds of lower-plex imaging followed by fluorophore dissociation or inactivation. For example, tissue movement (**Fig. 1m**) and tissue damage (**Fig. 1n**) caused cells present in early rounds of imaging to be lost at later cycles. These cells appear negative for all markers after movement or loss, confounding cell type assignment and leading to artefactual clusters in feature space (**Fig. 1o**). The extent of tissue loss varies between specimens and seems to arise during tissue dewaxing and antigen retrieval^31^ due to low tissue area (e.g., the TOPACIO fine-needle biopsies from patients 70, 89, 95, 96) and cellularity (e.g., adipose tissue).

The origins of some artefacts remain unknown, but likely arise from a combination of (i) pre-analytical variables—generally defined as variables arising prior to specimen staining, (ii) unwanted fluorescent objects (e.g., lint and antibody aggregates) introduced during staining, imaging, and washing steps, (iii) errors in data acquisition, and (iv) the intrinsic properties of the tissue itself^32,33^. The TOPACIO specimens (Dataset 1) were the most severely affected by these artefacts, whereas the CRC specimen (Dataset 2)^21^, which had been freshly-sectioned and carefully processed, was much less affected. However, only one slide was available from each TOPACIO patient, making repeat imaging impossible.

### Microscopy artefacts obscure analysis and interpretation of tissue-derived, single-cell data

Clustering Dataset 2 (CRC, CyCIF, ∼9.8×10^5^ total cells) with HDBSCAN yielded 22 clusters with 0.7% of cells remaining unclustered (**Fig. 2a**). Silhouette analysis^34^ showed that four clusters (6, 15, 17, and 21) remained under-clustered despite parameter tuning (**Fig. 2b**). Agglomerative hierarchical clustering of HDBSCAN clusters based on mean marker intensities revealed four meta-clusters (**Fig. 2c**) corresponding to tumor (meta-clusters A, B), stromal (C), and immune (D) cell populations. To study these 22 HDBSCAN clusters, cells from each cluster were selected at random and organized into galleries of 20 x 20 µm (30 x 30 pixel) image patches centered on reference nuclei (**Online Supplementary** Fig. 2). To facilitate interpretation, only the three most highly expressed protein markers were shown per cluster (based on channel intensities normalized across clusters; **Fig. 2c**). Inspection of these galleries showed that many clusters contained mixed cell types. For example, cluster 6 contained B cells, T cells, and stromal cells (**Fig. 2d**). The formation of clusters 9 and 11 were driven by bright antibody aggregates in the desmin and vimentin channels (**Fig. 2e, f**), respectively, whereas contaminating lint fibers led to the formation of cluster 12 (**Fig. 2g**). Cell loss was evident in cluster 14 (**Fig. 2h**), and cluster 10 comprised a domain of vimentin-positive tissue of unknown origin (**Fig. 2i**). Three additional clusters (2, 8, and 19; **Fig. 2j**) were caused by a region of tissue unexposed to antibodies during imaging cycle 3 as evidenced by a sharp cutoff in immunolabeling in this area. We reasoned that this artefact was likely due to human error during the performance of a complex 3D imaging experiment^21^. Clustering of Dataset 6 (CODEX, large intestine) also revealed clusters in which the expected separation of cell types was confounded by antibody aggregates, tissue folds, and image blur (Extended Data Fig. 2a-f and **Online Supplementary** Fig. 3). We conclude that the presence of image artefacts, even in relatively unaffected specimens, can drive formation of clusters that contain cells of different type (see **Supplementary Note 1** for a discussion of problems associated with background subtraction).

**Fig. 2.**
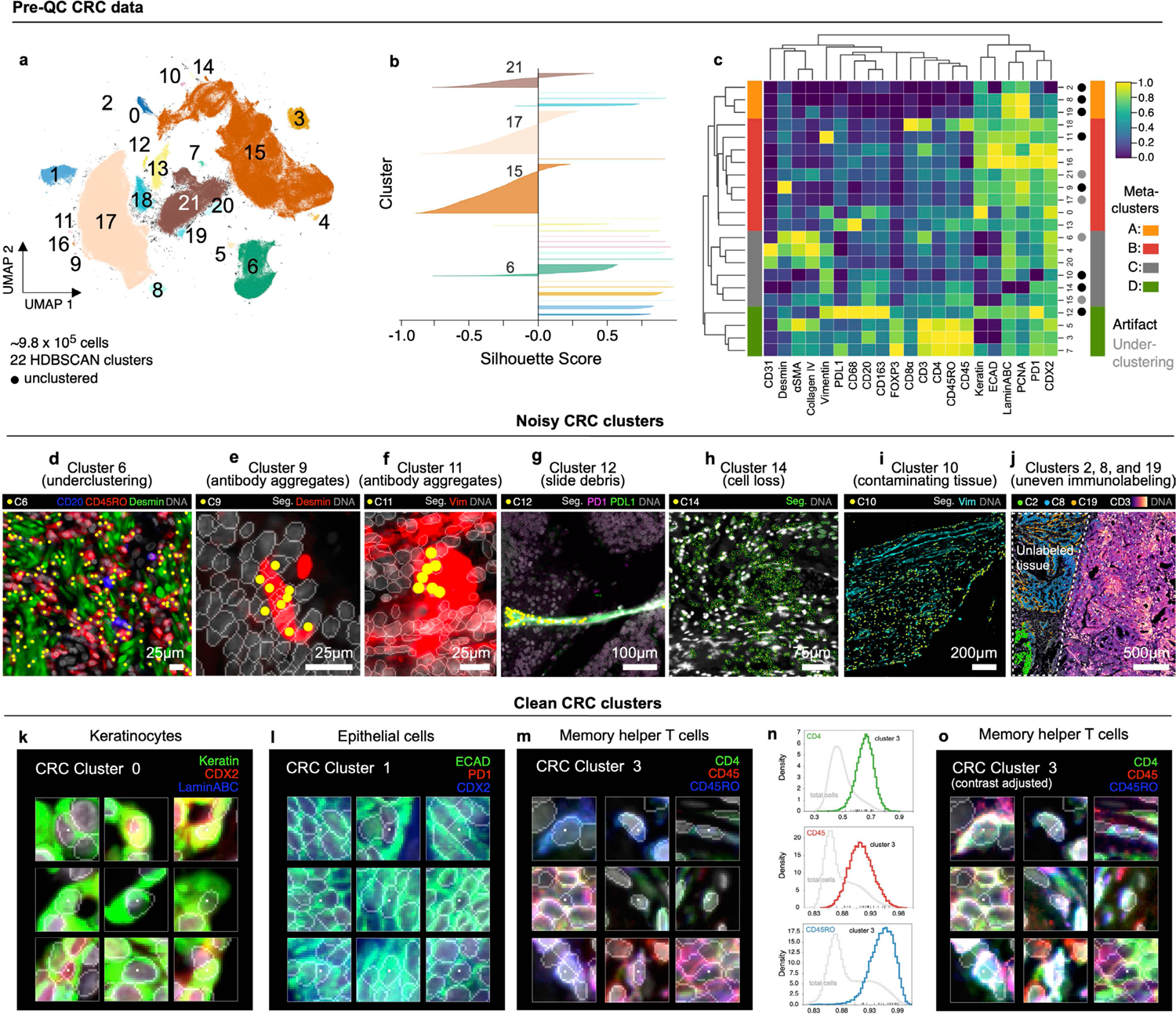
Evaluation of pre-QC cell clustering results from Dataset 2 (CRC). **a**, UMAP embedding of CRC data showing ∼9.8×10^5^ cells colored by HDBSCAN cluster (numbered 0-21). Black scatter points represent unclustered (ambiguous) cells. **b**, Silhouette scores for CRC clusters shown in panel (a). Clusters 6, 15, 17, and 21 exhibit cells with negative silhouette scores indicative of under-clustering. **c**, Clustered heatmap for CRC data showing mean signals of clustering cells normalized across clusters (row-wise). Four (4) meta-clusters defined by the heatmap dendrogram are highlighted. **d**, Cluster 6 cells (yellow dots) in a region of the CRC image demonstrating the co-clustering of distinct populations of B cells (CD20, blue), memory T cells (CD45RO, red), and stromal cells (desmin, green); Hoechst (gray) shown for reference. **e**, Anti-desmin antibody aggregates (red) in a region of the CRC image. Yellow dots highlight cluster 9 cells which have formed due to this artefact; Hoechst (gray) shown for reference. **f,** Anti-vimentin antibody aggregates (red) in a region of the CRC image. Yellow dots highlight cluster 11 cells that have formed due to this artefact; Hoechst (gray) shown for reference. **g**, Autofluorescent fiber in a region of the CRC image as seen in channels PD1 (magenta) and PD-L1 (green). Yellow dots highlight cluster 9 cells which have formed due to this artefact; Hoechst (gray) shown for reference. **h**, Cell loss in a region of the CRC image as indicated by anucleate segmentation outlines (green). Yellow dots highlight cluster 14 cells which have formed due to this artefact; Hoechst (gray) shown for reference. **i**, Contaminating (non-colonic) tissue at the top of the CRC image immunoreactive to anti-vimentin antibodies (cyan) comprising CRC cluster 10 (yellow dots); Hoechst (gray) shown for reference. **j**, Region of tissue at the bottom-left of the CRC image unexposed to antibodies during imaging cycle 3 which led to the formation of CRC clusters 2, 8, and 19; channels CD3 (colormap) and Hoechst (gray) shown for reference. **k-m**, Top three most highly expressed markers (1: green, 2: red, 3: blue) for clusters 0 (keratinocytes, **k**), 1 (crypt-forming mucosal epithelial cells, **l**), and 3 (memory helper T cells, **m**). A single white pixel at the center of each image patch highlights the reference cell. Nuclear segmentation outlines (translucent white outlines) and Hoechst (gray) shown for reference. **n**, Density histograms showing the distribution of cluster 3 cells according to channels CD4 (green outline), CD45 (red outline), and CD45RO (blue outline) superimposed on distributions of total cells according to the same channels (gray outlines). Rugplots at the bottom of each histogram show where 25 members of cluster 3 shown in panel (m) and Extended Data Fig. 2h reside in each distribution. **o**, Cluster 3 cells shown in panel (m) and Extended Data Fig. 2h after signal intensity cutoffs have been adjusted per image to improve the homogeneity of their appearance.

Many other clusters in Dataset 2 (e.g., 0, 1, 3, 7, and 16) contained few obvious artefacts. For example, cluster 0 comprised a phenotypically homogenous group of keratinocytes (**Fig. 2k**), while cluster 1 represented normal crypt-forming epithelial cells (**Fig. 2l**). Cluster 3 consisted of CD4, CD45, and CD45RO^+^ memory T cells distributed throughout the tissue (Extended Data Fig. 2g). Cells in this cluster appeared remarkably non-uniform (**Fig. 2m** and Extended Data Fig. 2h), despite their occupying a discrete region of the UMAP embedding (**Fig. 2a**) and having CD4, CD45, and CD45RO levels well above background (**Fig. 2n**). Protein expression among these cells was also well correlated (R=0.56 to 0.59; Extended Data Fig. 2i), suggesting that cluster 3 encompassed a single cell population. Consistent with this conclusion, adjusting image intensity on a per-channel and per-cell basis resulted in a more uniform appearance (**Fig. 2o** and Extended Data Fig. 2j,k). Cells in cluster 7 (Tregs, Extended Data Fig. 2l) also formed a tight cluster (**Fig. 2a**) with good correlation in expression of CD4, CD45, and CD45RO (R=0.51 to 0.62; Extended Data Fig. 2m) but weak correlation with FOXP3, the defining transcription factor for Tregs (R=0.13 to 0.31; Extended Data Fig. 2n). We conclude that nonuniformity in the appearance of these cells likely arises from natural cell-to-cell variation in protein levels^35^— not simply dataset noise—but that multidimensional clustering correctly groups such cells into biologically meaningful subtypes. Thus, visual review must be performed with care, and ideally in conjunction with data-driven approaches such as HDBSCAN.

Clustering Dataset 1 (25 TOPACIO specimens) gave rise to 492 HDBSCAN clusters with ∼29% of cells remaining unclustered (**Fig. 3a**) and exhibiting no discernible spatial pattern in the underlying images (Extended Data Fig. 3a). Most clusters were associated with positive silhouette scores, indicating a good fit (**Fig. 3b**). While a few small clusters contained cells from a single tissue specimen (e.g., cluster 75 with 418 cells and cluster 146 with 2140 cells), most clusters (441/492) contained cells from more than half of the 25 TOPACIO specimens (Extended Data Fig. 3b); nevertheless, even these clusters often contained fewer than 3,000 cells (**Fig. 3c**). Agglomerative hierarchical clustering generated six meta-clusters (**Fig. 3d**), but the heatmap revealed an unusual dichotomy of very bright signals for some markers and dim signals for others. Only meta-cluster C, which comprised 57% of the cells exhibited graded signals across all channels (**Fig. 3d,e**). Image patches from a random set of 48 clusters revealed the presence of numerous tissue and imaging artefacts, including bright fluorescent signals, over-saturated nuclear stains, and poor segmentation (**Fig. 3f-h** and **Online Supplementary** Fig. 4). Cluster 15 (meta-cluster A) arose from an image alignment error at the bottom of TOPACIO specimen 55 (Extended Data Fig. 3c) and meta-clusters B, D, E, and F were caused by the presence of cells with channel intensities at or near zero as a result of image background subtraction (see **Supplementary Note 1** and **Supplementary** Fig. 2).

**Fig. 3.**
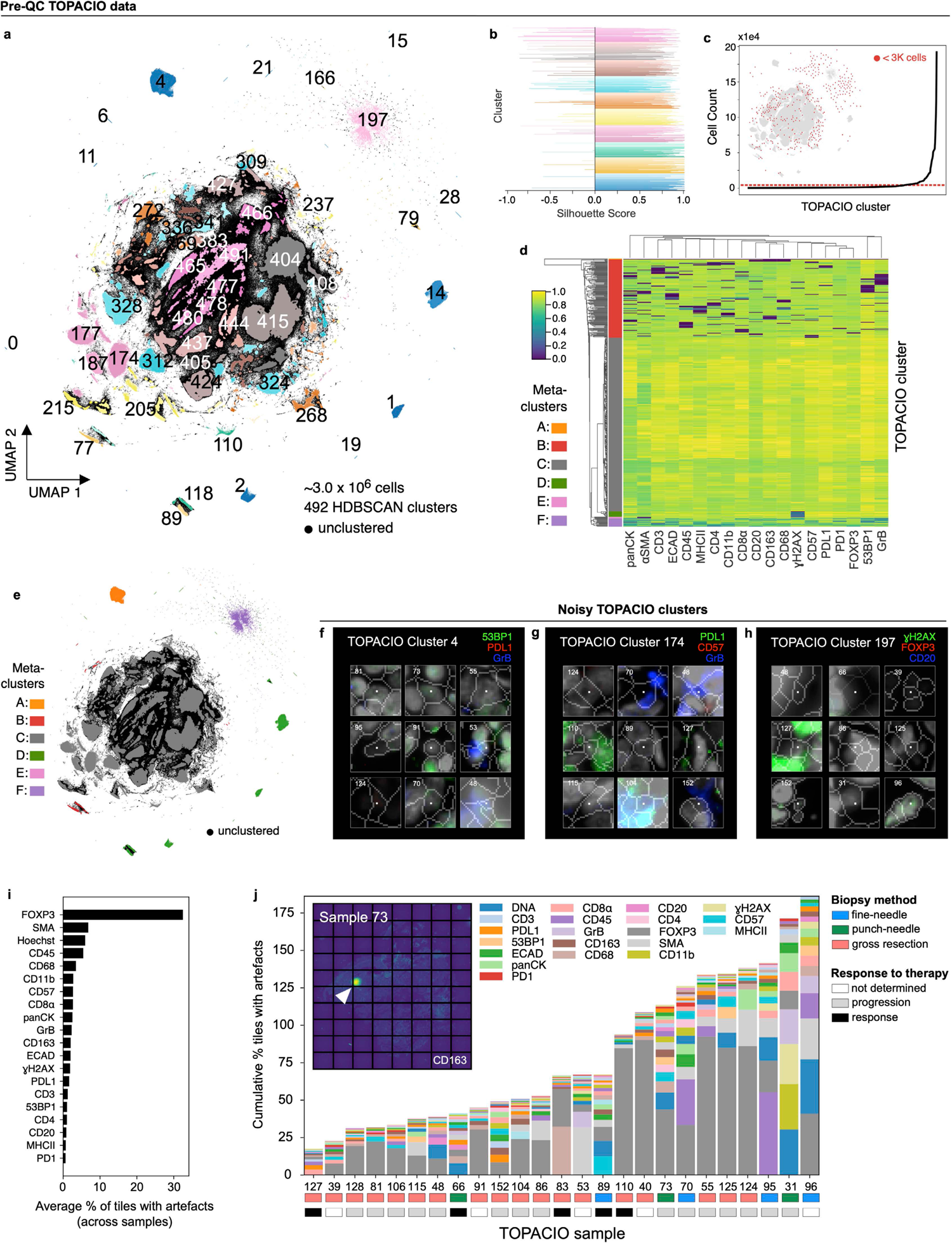
Evaluation of pre-QC cell clustering results from Dataset 1 (TOPACIO). **a,** UMAP embedding of ∼3×10^6^ cells from the TOPACIO dataset colored by HDBSCAN cluster. Black scatter points represent unclustered (ambiguous) cells. **b**, Silhouette scores for TOPACIO clusters shown in panel (a). **c**, Line plot showing cell counts per TOPACIO cluster. Clusters with cell counts below the horizonal dashed red line are those with fewer than 3K cells which are highlighted in the TOPACIO embedding (inset) by red scatter points at their relative positions. **d**, Clustered heatmap of clusters from TOPACIO data showing mean signal intensities of clustering cells normalized across clusters (row-wise). Six (6) meta-clusters defined by the heatmap dendrogram at the left are highlighted. **e**, TOPACIO embedding colored by meta-clusters shown in panel (d). **f-h**, Top three most highly expressed markers (1: green, 2: red, 3: blue) for TOPACIO clusters 4 (**f**), 174 (**g**), and 197 (**h**) which were all severely affected by dataset noise. A single white pixel at the center of each image highlights the reference cell. Nuclear segmentation outlines (translucent white outlines) and Hoechst (gray) are shown for reference. **i**, Bar chart showing the average percentage of image tiles affected by a visual artefact across the 25 TOPACIO specimens; marker identities at left denote the affected channel. **j**, Stacked bar chart showing the cumulative percentage of channel-specific image tiles per TOPACIO specimen affected by miscellaneous visual artefacts. Because these artefacts can impact multiple channels at the same time, cumulative percentages can be higher than 100%. Inset shows an example illumination aberration in the CD163 channel of TOPACIO specimen 73. Categories for tissue biopsy method and patient treatment response are indicated below each bar. Artefacts were found to be less abundant in tissue resections as compared to fine-needle and punch-needle biopsies as determined by one-way ANOVA followed by pairwise Tukey’s HSD (F = 10.27, p = 0.0007; fine-needle vs. resection mean difference = 204.83, p-adj = 0.0145; resection vs. punch-needle mean difference = −283.0, p-adj: 0.0029).

To estimate the prevalence of visible artefacts in Dataset 1, we generated a set of down-sampled single-channel images with tile gridlines superimposed and manually estimated the number of tiles impacted by overt artefacts (**Online Supplementary** Fig. 5). This showed that ∼5,490 of 156,300 tiles (3.5%) were affected by antibody aggregates, folds, illumination aberrations, or slide debris. The FOXP3 channel was the worst affected (>30% of tiles; **Fig. 3i**) involving streaks of non-specific antibody signal. Artefacts were less abundant in tissue resections as compared to fine-needle and punch-needle biopsies (one-way ANOVA, Tukey’s HSD: p-adj = 0.0029 to 0.0145) but there was no correlation with response to therapy (F = 0.40, p = 0.67, **Fig. 3j**). We concluded that the presence of imaging artefacts causes single-cell analysis methods to fail with TOPACIO data, but that errors were not preferentially biased with respect to patient response.

### Identifying and removing noisy single-cell data with CyLinter

To remove imaging artefacts from tissue images via computer-assisted human review, we developed CyLinter as a plugin for the Napari^23^ multi-channel image viewer (**Fig. 4a** and Extended Data Fig. 4).

**Fig. 4.**
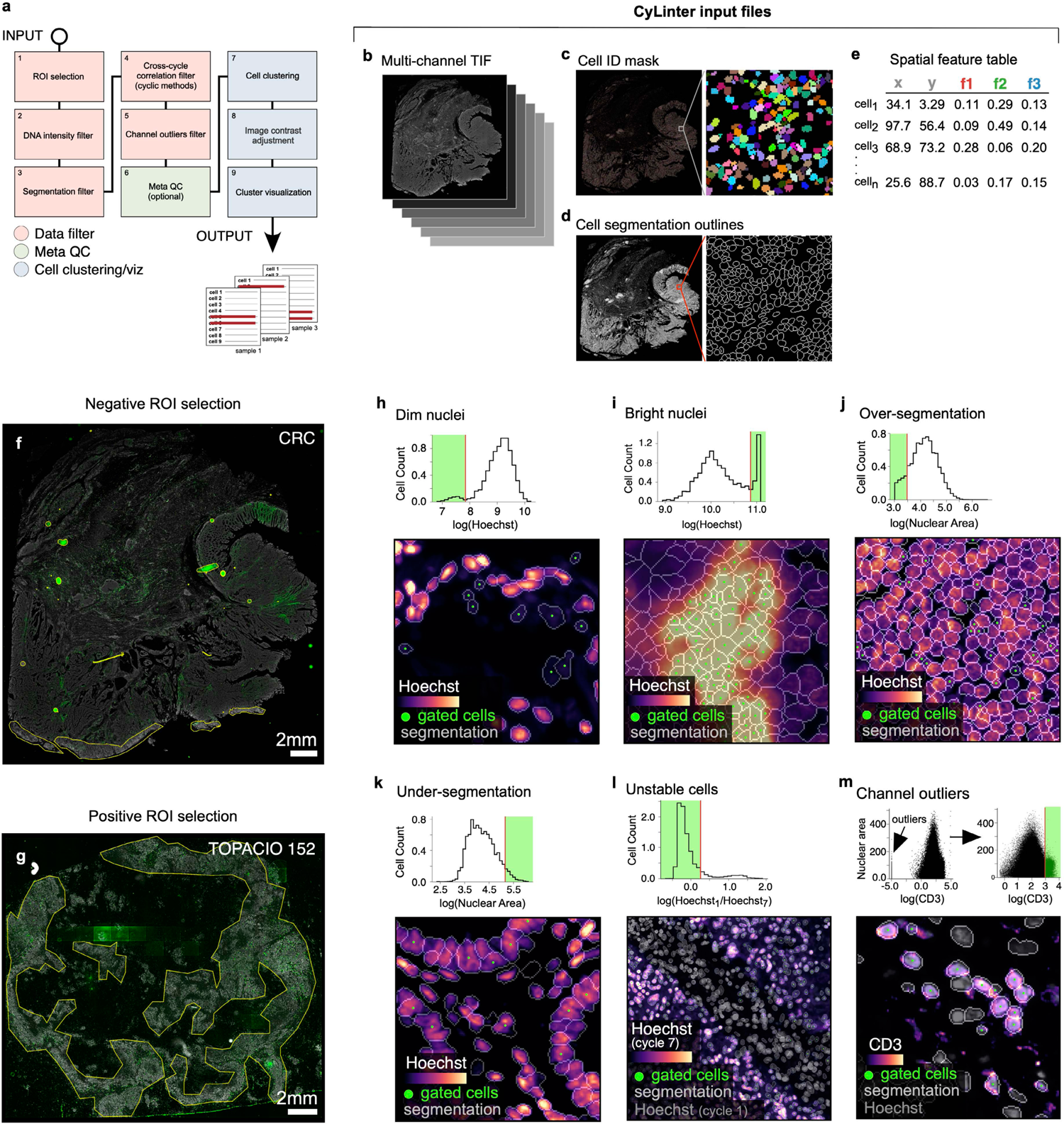
Identifying and removing noisy single-cell data points with CyLinter. **a**, Schematic representation of the CyLinter workflow. Modules are colored by type: data filtration (red), metaQC (green), cell clustering/visualization (blue). **b-e**, CyLinter input: **b**, Multiplex image file, **c**, Cell ID mask, **d**, Cell segmentation outlines, **e**, Single-cell feature table. **f**, Negative ROI selection in CyLinter. Dataset 2 (CRC) is shown with ROIs (yellow outlines) applied to various artefacts in the CD163 channel which will be dropped from subsequent analysis. **g**, Positive ROI selection in CyLinter. Dataset 1 (TOPACIO, specimen 152) is shown with ROIs (yellow outlines) applied to regions devoid of artefacts in the FOXP3 channel which will be retained for further analysis. **h**, Filtering dim nuclei. Top: Density histogram of mean Hoechst signal for cells in Dataset 3 (EMIT TMA, core 12, non-neoplastic lung). Bottom: Hoechst (colormap) in a region of the same core demonstrating dim nuclei (green dots) falling to the left of the red gate in the corresponding histogram. Nuclear segmentation outlines are shown for reference (translucent outlines). **i**, Filtering bright nuclei. Top: Density histogram of mean Hoechst signal for Dataset 1(TOPACIO, specimen 110). Bottom: Hoechst (colormap) in a region of the same specimen demonstrating bright nuclei (green dots) caused by tissue bunching that fall to the right of the gate in the corresponding histogram. Nuclear segmentation outlines are shown for reference (translucent outlines). **j**, Filtering over-segmented cells. Top: Density histogram of mean Hoechst signal for Dataset 2 (CRC). Bottom: Hoechst (colormap) in a region of the specimen demonstrating over-segmented cells (green dots) falling to the left of the red gate in the corresponding histogram. Nuclear segmentation outlines are shown for reference (translucent outlines). **k**, Filtering under-segmented cells. Top: Density histogram of mean Hoechst signal for Dataset 3 (EMIT TMA, core 84, non-neoplastic colon). Bottom: Hoechst (colormap) in a region of the specimen demonstrating under-segmented cells (green dots) falling to the right of the red gate in the corresponding histogram. Nuclear segmentation outlines are shown for reference (translucent outlines). **l**, Filtering unstable cells. Top: Density histogram of the log(ratio) between Hoechst signals from the first and last CyCIF imaging cycles for Dataset 3 (EMIT TMA, core 74, renal cell carcinoma). Bottom: Hoechst (last cycle, colormap) superimposed on Hoechst (first cycle, gray) in a region of the specimen demonstrating the selection of stable cells (green dots) falling to the left of the red gate in the corresponding histogram. Nuclear segmentation outlines are shown for reference (translucent outlines). Note: unlike panels (h-k) which highlight cells that will be excluded from an analysis, cells highlighted in this panel will be retained for further analysis. **m**, Filtering channel outliers. Top: Scatter plot showing CD3 (x-axis) vs. nuclear segmentation area (y-axis) of cells from Dataset 1 (TOPACIO, specimen 152) before (left) and after (right) outlier removal and signal rescaling (0-1). Bottom: CD3 (colormap) and Hoechst (gray) signals in a region of the same specimen with CD3^+^ cells (green dots) falling to the right of the red gate in the scatter plot in which outliers have been removed. Nuclear segmentation outlines are shown for reference (translucent outlines).

CyLinter consists of a set of QC software modules written in the Python programming language that process images and corresponding single-cell data in a flexible manner in which modules can be run iteratively while bookmarking progress within and between modules. CyLinter takes four files as input for each tissue specimen: 1) a stitched and registered multiplex image (TIFF/OME-TIF), 2) a cell identification mask generated by a segmentation algorithm, 3) a binary image showing the boundaries between segmented cells, and 4) a spatial feature table^19^ in CSV format comprising the location and computed signal intensities for each segmented cell (**Fig. 4b-e**, respectively). With a dataset comprising multiple images and spatial feature tables, CyLinter automatically aggregates the data into a single Pandas (Python) dataframe^36^ for efficient processing (Extended Data Fig. 4a). CyLinter then removes of artefactual cells from the dataframe (see https://labsyspharm.github.io/cylinter/ for implementation details) with no attempt to infer missing values.

The first CyLinter module, *selectROIs* (Extended Data Fig. 4b), lets the user view a multi-channel image and manually identify artefacts such as regions of tissue damage, antibody aggregates, and large illumination aberrations. Lasso tools native to the Napari image viewer are used to define regions of interest (ROIs) corresponding to artefacts. We found that negative selection (in which highlighted cells are dropped from further analysis) worked effectively for Dataset 2 (CRC, **Fig. 4f**), but Dataset 1 (TOPACIO) was affected by too many artefacts for this approach to be efficient. Thus, CyLinter implements an optional positive ROI selection mode, in which users select tissue regions devoid of artefacts for retention in the dataset (**Fig. 4g**).

CyLinter also includes an automated companion algorithm that works with the *selectROIs* module to programmatically flag likely artefacts for human review (Extended Data Fig. 4b and **Methods**). This efficiently identifies features with intensities outside the distribution of biological signals.

CyLinter’s *dnaIntensity* module (Extended Data Fig. 4c) allows users to inspect histogram distributions of per-cell mean nuclear intensities. Nuclei at the extreme left side of the distribution often correspond to cells lying outside of the focal plane (**Fig. 4h**) and those to the right side correspond to cells oversaturated with DNA dye or found in tissue folds (**Fig. 4i**). This module redacts data based on lower and upper thresholds initially defined by Gaussian Mixture Models (GMMs) that can be manually refinement if necessary. Instances of substantial over and under-segmentation can be identified based on the area of each segmentation instance followed by removal using the *dnaArea* module (Extended Data Fig. 4d). This method was particularly effective at removing many over-segmented cells in the CRC image (**Fig. 4j**) and under-segmented cells frequently encountered among tightly-packed columnar epithelial cells in normal colon specimens (e.g., EMIT TMA core 84; **Fig. 4k**).

In cyclic imaging methods, nuclei are re-imaged every cycle and individual cells are sometimes lost due to tissue movement or degradation^37,38^. CyLinter’s *cycleCorrelation* module (Extended Data Fig. 4e) computes histograms of log_10_-transformed DNA intensity ratios between the first and last imaging cycles (log_10_[DNA_1_/DNA_n_]); cells that remain stable give rise to ratios around zero, whereas those that are lost give rise to a discrete peak with ratios > 0. Gating the resulting histogram on stable cells eliminates unstable cells from the data table (**Fig. 4l**). Protein signals are then log-transformed (Extended Data Fig. 4f). The *pruneOutliers* module makes it possible to visualize scatter plots of per-cell signals from all specimens in a multi-image dataset and remove residual artefacts (e.g., small antibody aggregates) based on lower and upper percentile cutoffs (**Fig. 4m** and Extended Data Fig. 4g). Cells falling outside of the thresholds can be visualized to ensure that selected data points are indeed artefacts.

The *dnaIntensity*, *dnaArea*, *cycleCorrelation* and *pruneOutliers* modules all provide a linked view of the original image in which cells to be included or excluded by the user’s chosen threshold settings are directly overlaid for visual confirmation of threshold accuracy. These labels are dynamically updated as the thresholds are adjusted. This “visual review” is crucial to ensuring that true cell populations that happen to have extreme variations in size or signal intensity are not accidentally removed.

### Correcting for bias in user-guided histology QC via unsupervised cell clustering

Human-guided artefact detection is subject to errors and biases and the *metaQC* module (Extended Data Fig. 4h) addresses this by performing unsupervised clustering on equal size combinations of redacted and retained data. Cells flagged for redaction that fall within predominantly clean clusters in retained data can be added back to the dataset, while those retained in the dataset that co-cluster with predominantly noisy cells (presumed to have been missed during QC) can be removed from the data table. The *PCA* module (Extended Data Fig. 4i) performs Horn’s parallel analysis to help the user determine whether 2 or 3 principal components should be used in the *clustering* module (described below). The *setContrast* module (Extended Data Fig. 4j) allows users to adjust per-channel image contrast on a reference image and then apply these settings to all images in a batch. Like the *metaQC* module, CyLinter’s *clustering* module (Extended Data Fig. 4k) allows users to perform UMAP^39^ or t-SNE^40^ data dimensionality reduction and HDBSCAN^29^ density-based clustering to identify discrete cell populations in high-dimensional feature space; the *clustermap* module (Extended Data Fig. 4l) generates high-dimensional protein expression profiles for each cluster. To test for statistical differences in cell type frequency between tissues associated with test and control conditions (e.g., treated vs. untreated) the *sampleMetadata* field in CyLinter’s configuration file can be populated and the *frequencyStats* module (Extended Data Fig. 4m) can be run. The *curateThumbnails* module (Extended Data Fig. 4n) automatically draws cells at random from each identified cluster and generates image galleries for efficient visual inspection. Together, these QC steps allow a user to apply a series of objective criteria to redacted and retained data to revise the output of the prior data filtration modules. On completion of the QC pipeline, CyLinter returns a single redacted spatial feature table together with a QC report for reproducibly and transparency of the analysis. Artefacts identified by CyLinter are ideal for training machine learning models that can automate artefact detection; we have therefore created a public repository for CyLinter QC reports and artefact libraries (see **Supplementary Note 2** and **Supplementary** Fig. 3).

### Impact of CyLinter-based quality control on the CRC and TOPACIO datasets

Applying CyLinter to Dataset 2 (CRC) resulted in the removal of ∼23% of total cells (**Fig. 5a**). Over-segmentation was the largest problem, affecting ∼16% of cells (Extended Data Fig. 5a), with ∼4% or less dropped by the other QC modules. Thus, better segmentation would in principle have allowed ∼93% of the data to be retained. Using HDBSCAN in CyLinter’s *clustering* module, we identified 78 clusters (**Fig. 5b**), 56 more than pre-QC data (**Fig. 2a**). Silhouette scores were predominantly positive, suggesting effective clustering (**Fig. 5c**). Agglomerative hierarchical clustering yielded six meta-clusters with marker expression patterns corresponding to populations of tumor cells (meta-cluster A; **Fig. 5d**), stromal cells (B), memory T cells (C), macrophages (D), B cells (E), and effector T cells (F). Using the *curateThumbnails* module, we confirmed that all 78 clusters were largely free of visual artefacts (**Fig. 5e-g** and **Online Supplementary** Fig. 6). The increase in the number of clusters in the post-QC CRC embedding appeared to be due to the removal of pre-QC outliers that constrained the remainder of cells to a relatively narrow region of UMAP feature space. For example, by coloring the pre-QC embedding by post-QC CRC clusters, we found that pre-QC cluster 6 (**Fig. 2a-d**) consisted of nine different cell populations in the post-QC embedding (**Fig. 5h-j**). These included vimentin^+^ mesenchymal cells (post-QC cluster 9), memory CD8^+^ T cells (post-QC cluster 51), and collagen IV^+^ stromal cells (post-QC cluster 54). Similar analyses performed on Dataset 6 (CODEX) showed comparable improvements in the post-QC UMAP embedding, HDBSCAN clustering, and associated heatmap of cluster protein expression profiles (Extended Data Fig. 5b-h and **Online Supplementary** Fig. 7). We conclude that post-QC clusters represent *bona fide* cell states that are better distributed across biologically meaningful regions of the UMAP embedding.

**Fig. 5.**
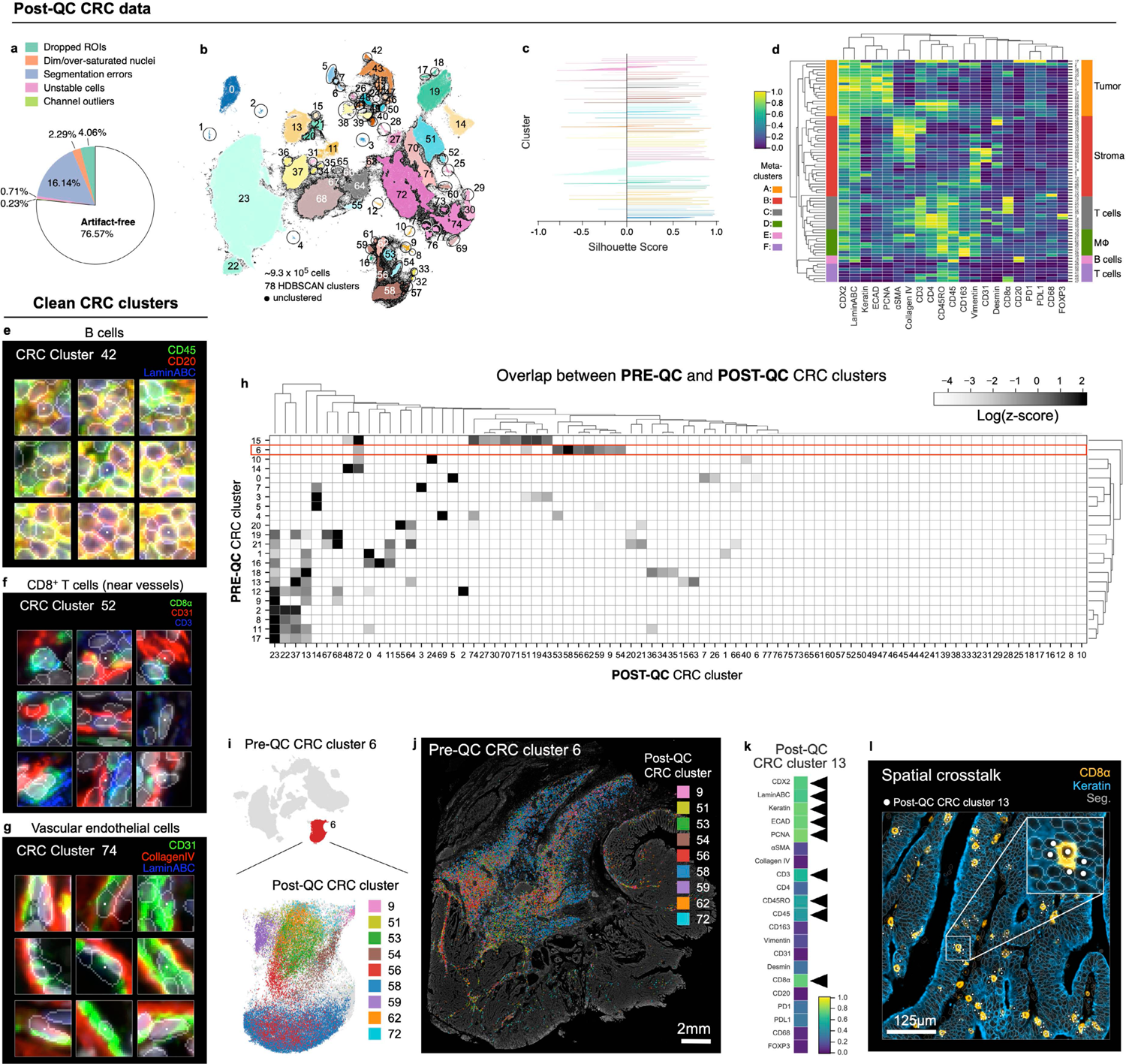
Cleaning Dataset 2 (CRC) with CyLinter. **a**, Fraction of cells in Dataset 2 redacted by each QC filter in the CyLinter pipeline. Dropped ROIs = cells dropped by *selectROIs* module), Dim/over-saturated nuclei = cells dropped by *dnaIntensity* module, Segmentation errors = cells dropped by *areaFilter* module, Unstable cells = cells dropped by *cycleCorrelation* module, Channel outliers = cells dropped by *pruneOutliers* module, Artefact-free = cells remaining after QC. **b**, UMAP embedding of post-QC CRC data showing ∼9.3×10^5^ cells colored by HDBSCAN cluster. Black scatter points represent unclustered (ambiguous) cells. **c**, Silhouette scores for post-QC CRC clusters shown in panel (b), **d**, Clustered heatmap of post-QC CRC clusters showing mean signal intensities of clustered cells normalized across clusters (row-wise). Six (6) meta-clusters defined by the clustered heatmap dendrogram at the left are highlighted. **e-g**, Top three most highly expressed markers (1: green, 2: red, 3: blue) for post-QC CRC clusters 42 (B cells, **e**), 52 (CD8^+^ T cells near blood vessels— formed as a side effect of spatial crosstalk, **f**), and 74 (vascular endothelial cells, **g**). A single white pixel at the center of each image highlights the reference cell. Nuclear segmentation outlines (translucent outlines) and Hoechst (gray) shown for reference. **h**, Overlap between pre-QC CRC clusters (rows) and post-QC CRC clusters (columns) showing pre- and post-QC clusters have a one-to-many correspondence. **i**, Pre-QC CRC embedding showing the position of cluster 6 (red, inset) and its composition according to post-QC CRC clusters. **j**, Locations of cells in pre-QC cluster 6 colored by their post-QC cluster label showing that pre-QC cluster 6 is composed of cells occupying distinct regions throughout the muscularis propria of the CRC image—a non-cancerous, smooth muscle-rich region of tissue. **k**, Mean signal intensities for post-QC CRC cluster 13 cells. Black arrows point to bright channels consistent with both epithelial cells and CD8^+^ T cells. **l**, Post-QC CRC cluster 13 cells (white dots) shown in context of the CRC image demonstrating spatial crosstalk between keratin^+^ tumor cells (blue) and CD8^+^ T cells (orange). Nuclear segmentation outlines (translucent outlines) shown for reference.

Despite improvements in post-QC clustering of Dataset 2 (CRC), visual inspection of the clustered heatmap (**Fig. 5d**) continued to reveal cells with unexpected marker expression patterns. For example, post-QC cluster 13 contained cells with epithelial markers such as Keratin and ECAD and T cell markers such as CD3, CD45RO, CD45, and CD8α (**Fig. 5k**). There is no known cell type that expresses this marker combination.

Visual inspection showed that cluster 13 consisted of CD8^+^ T cells surrounded by keratin positive tumor cells (**Fig. 5l**). Because segmentation is not perfect, pixels from CD8^+^ T cells were incorrectly assigned to neighboring epithelial cells and *vice versa*, a phenomenon known as spatial crosstalk (or lateral spillover)^41^. Tools such as REDSEA^41^ attempt to address this problem, but instances of crosstalk must currently be identified in post-QC data through inspection of heatmaps and cell image galleries.

In the case of Dataset 1 (TOPACIO), CyLinter removed 84% of cells, with most (∼53%) removed during positive ROI selection (**Fig. 6a**). Bright outliers primarily attributed to antibody aggregates (∼14% of cells), cell detachment with increasing cycle number (12%), segmentation errors (4%), and dim/over-saturated nuclei (1%) were also common in this dataset. Cells redacted by CyLinter for both the CRC and TOPACIO datasets exhibited no discernable pattern in spatial location (Extended Data Fig. 6a,b) and data redacted from the TOPACIO specimens was not biased with respect to biopsy type (one-way ANOVA, F = 1.93, p = 0.17) or treatment response (F = 0.71, p = 0.50). Overall, the post-QC TOPACIO dataset comprised 43 clusters among ∼3.0×10^6^ cells (**Fig. 6b**). Silhouette analysis revealed positive scores for all clusters except 42 which represented the majority of tumor cells in these specimens (**Fig. 6c**). We found that tumor cell populations tended to cluster by patient, whereas immune cell populations tended to be more heterogenous with respect to patient ID (Extended Data Fig. 6c). Agglomerative hierarchical clustering based on mean marker intensities yielded four meta-clusters corresponding to stromal (meta-cluster A; **Fig. 6d**), tumor (B), lymphoid (C), and myeloid (D) cells. CyLinter’s *curateThumbnails* module revealed that most cells had a high degree of concordance in morphology and marker expression and were consistent with known cell types (**Fig. 6e-i** and **Online Supplementary** Fig. 8). For example, post-QC TOPACIO cluster 0 corresponded to cells with small, round, nuclei with intense plasma membrane staining for CD4 and nuclear staining for FOXP3, consistent with T regulatory cells (Tregs, **Fig. 6e**), cells in cluster 21 were high in panCK and γH2AX, indicative of breast cancer cells containing DNA damage (**Fig. 6g**), and cells in cluster 35 were conventional CD4^+^ helper T cells adjacent to panCK^+^ tumor cells (captured as a manifestation of spatial crosstalk; **Fig. 6h**). Like in Dataset 2 (CRC), by coloring the post-QC embedding by pre-QC cluster labels, we found that many pre-QC clusters were composed of different post-QC cell types (**Fig. 6j**). For example, pre-QC cluster 404 consisted of CD8^+^ T cells (which mapped to post-QC cluster 5), CD4^+^ T cells (post-QC cluster 10), αSMA^+^ stromal cells (post-QC cluster 24), and CD68^+^ macrophages (post-QC cluster 39). Thus, imaging artefacts in the TOPACIO data not only resulted in an unrealistically large number of clusters, but these clusters still contained mixed cell types.

**Fig. 6.**
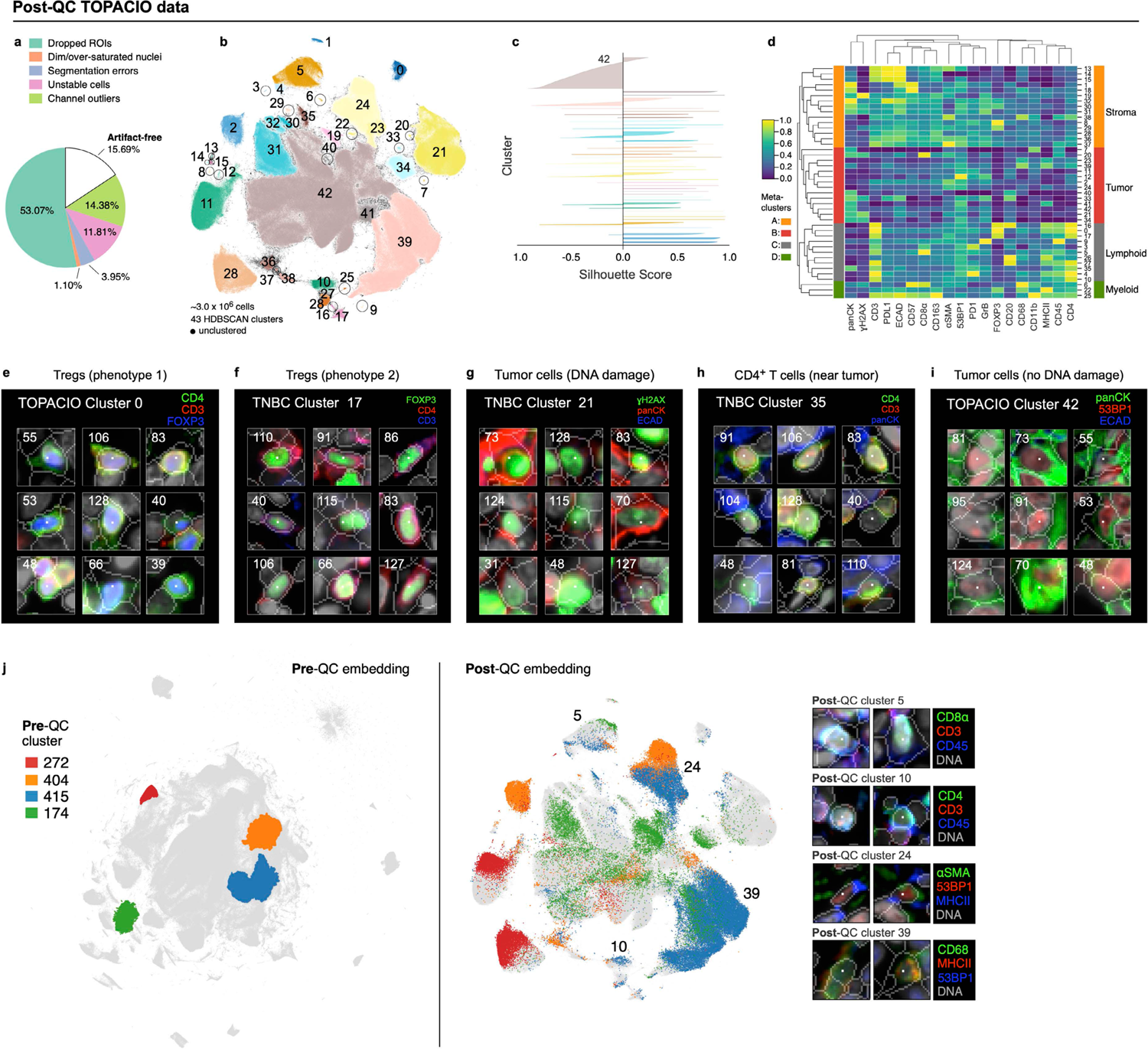
Cleaning Dataset 1 (TOPACIO) with CyLinter. **a**, Fraction of cells in the TOPACIO dataset redacted by each QC filter in the CyLinter pipeline. Dropped ROIs = cells dropped by *selectROIs* module, Dim/over-saturated nuclei = cells dropped by *dnaIntensity* module, Segmentation errors = cells dropped by *areaFilter* module, Unstable cells = cells dropped by *cycleCorrelation* module, Channel outliers = cells dropped by *pruneOutliers* module, Artefact-free = cells remaining after QC. **b**, UMAP embedding of TOPACIO data showing ∼3.0×10^6^ cells colored by HDBSCAN cluster. Black scatter points represent unclustered (ambiguous) cells. **c**, Silhouette scores for post-QC TOPACIO clusters shown in panel (b). Cluster 42 is an under-clustered population. **d**, Clustered heatmap for clusters from post-QC TOPACIO data showing mean signal intensities of clustered cells normalized across clusters (row-wise). Four (4) meta-clusters defined by the clustered heatmap dendrogram at the left are highlighted. **e-i**, Top three most highly expressed markers (1: green, 2: red, 3: blue) for clusters 0 (Tregs: phenotype 1, **e**), 17 (Tregs: phenotype 2, **f**), 21 (breast cancer cells with DNA damage, **g**), 35 (CD4^+^ T cells near breast cancer cells, **h**), and 42 (breast cancer cells without DNA damage, **i**). A single white pixel at the center of each image highlights the reference cell. Nuclear segmentation outlines (translucent outlines) and Hoechst (gray) shown for reference. **j**, Left: Pre-QC TOPACIO UMAP embedding (also shown in **Fig. 3a**) with the location of five clusters selected and highlighted at random. Right: Location of the cells from the four pre-QC clusters shown in the embedding at left in the context of the post-QC TOPACIO UMAP embedding (also shown in panel b) demonstrating that these pre-QC clusters actually consisted of different cell types. Image patches of cells representing post-QC clusters are shown at far right.

## DISCUSSION

In this paper we show that artefacts commonly present in highly multiplexed tissue images have a dramatic impact on single-cell analysis. These artefacts can be broadly subdivided into: (i) those intrinsic to the specimen itself such as tissue folds and hair or lint, (ii) those arising during staining and image acquisition such as antibody aggregates, and (iii) those arising during image-processing such as cell segmentation errors. The first class is unavoidable and does not usually interfere with visual review by human experts. The second and third classes can be minimized but not fully eliminated by good experimental practices. However, even relatively infrequent artefacts as in datasets 2 (CyCIF) and 6 (CODEX) can strongly impact clustering and other types of single cell analysis. Archival specimens stored in paraffin blocks or mounted on slides years prior to imaging and are even more problematic insofar as artefacts are common and only one slide may be available for each specimen; unfortunately, this is not unusual in correlative studies of completed clinical trials.

The presence of cells affected by imaging artefacts has complex effects on clustering algorithms used to identify cell types and states. It can generate large numbers of spurious clusters but also cause these clusters to contain cells of multiple types. Removing the problematic cells using CyLinter solves this problem. When data are removed, there is always concern that findings will be biased. CyLinter addresses this in several ways, including by visual review of filtered cells against the image itself, performing meta-analysis of redacted features (*metaQC*), performing specimen subgroup analysis, and by generating a QC report for each specimen or set of specimens; the latter should ideally be included with all datasets. Similar issues arise with single cell sequencing, although much of the problem occurs during tissue dissociation, microfluidic or flow cytometry sorting, and library preparation^42,43^. An advantage of tissue imaging is that redacted data can be inspected in the context of the original image to identify patterns indicative of selection bias.

Quality control is recognized as a critical step in the acquisition of scRNA-Seq data and a robust ecosystem of QC tools has therefore been developed^42,44^. In contrast, CyLinter is among the first tools for QC of highly multiplexed tissue images. CyLinter is designed to accelerate and systematize human visual review, making it compatible with a wide range of tissue types. Efficiency is increased through automated ROI curation, smart thresholding using GMMs, and use of multi-specimen dataframes. We found that even the badly affected set of 25 specimens representing the TOPACIO dataset took a single reviewer less than a week to clean, which compares favorably with several weeks needed collect the data and several months or more to perform detailed spatial analysis. More automated approaches would nonetheless be valuable, and in **Supplementary Note 2** we describe a proof-of-concept DL model for artefact identification. The area under the receiver operator curve (ROC) of ∼0.73 shows that the approach is feasible, but that performance is not yet adequate for general use. It seems highly likely that this reflects insufficient and insufficiently diverse training data. CyLinter is the ideal way to generate this training data and we have therefore created a public artefact repository linked to the CyLinter website to collect data that can be used for progressive improvement of our DL model or models developed by others.

Microscopy is traditionally a visual field and our experience with over 1,000 whole-slide high-plex images from dozens of tissue and tumor types has demonstrated that spatial feature tables generated using existing algorithms not only contain errors and omissions, but they also poorly represent much of the morphological information in images. This emphasizes the necessity of visual review: any hypothesis generated through analysis of data in a spatial feature table must be confirmed through inspection of the underlying images. At the same time, visual review must be backed up by objective methods that detect and correct for human errors and biases. The QC tools in CyLinter achieve this combination of human review and algorithmic backup and represent one key step in making single cell analysis of high-plex spatial profiles more interpretable and reproducible.

### Supplementary Note 1: Impact of image background subtraction on derived single-cell data

Background subtraction is commonly used with multiplexed imaging to remove autofluorescence and fluorescence arising from non-specific antibody binding to the specimen. However, we identified a number of challenges associated with this approach. For example, plotting histograms of the distribution of per-cell signal intensities channel in the pre-QC TOPACIO dataset revealed small numbers of cells with zero-valued signal intensities in all channels (**Supplementary** Fig. 2a). We reasoned that this effect was due to rolling ball image background subtraction^45^ which was used to increase antibody signal-to-noise, but which had the unanticipated consequence of creating cells with signal intensities equal to zero that, after log-transformation, were far lower than values associated with other cells in the image. This effect was readily observed when the UMAP embedding was colored by channel signal intensity, as it revealed small clusters of extremely dim cells among much larger numbers of clusters whose signals were comparatively bright (**Supplementary** Fig. 2b,c). Using the panCK channel to better understand how cells with low signal intensities impacted the TOPACIO clustering result, we found that clusters within meta-cluster B (e.g., cluster 14) were exclusively composed of cells with zero-valued signals, while those in meta-cluster C (e.g., cluster 174) had signals that were all > 0, and those in meta-cluster F (e.g., cluster 197) were comprised of a mixture of cells with zero and non-zero signals (**Supplementary** Fig. 2d). The simple removal of cells with zero-value signal intensities from the pre-QC TOPACIO dataset (with no other quality control measures) eliminated small dark clusters characterized by very low signal intensities and significantly increased the resolution between immunopositive and immunonegative cell populations as seen in both the channel intensity histograms (**Supplementary** Fig. 2e) and UMAP embeddings colored by channel (**Supplementary** Fig. 2f). Resolution between positive and negative cells was further improved in the post-QC TOPACIO clustering after the removal of cells with near-zero signal intensities in addition to other artefacts (**Supplementary** Fig. 2g,h). This was also true of Dataset 6 (CODEX; **Supplementary** Fig. 2i,j). Thus, while background subtraction is useful for improving data quality, especially for low signal-to-noise antibodies, our analysis shows that it can skew the natural distribution of protein signals in an image and have a profound effect on the interpretation of single-cell data due to the spurious formation of irrelevant cell clusters. When using background subtraction, it is important to control for these problems.

### Supplementary Note 2: Developing a DL model for automated artefact detection

Although tools based on visual review are common in microscopy, there are obvious benefits to machine learning approaches^46–49^. To generate initial training data for a DL model to automatically flag arbitrary artefacts in multiplex IF images, three human annotators assembled ground truth artefact masks for 24 CyCIF channels in 11 serial tissue sections of the CRC dataset analyzed in this study (Dataset 2, **Supplementary** Fig. 1b). Single channel images (and their corresponding ground truth artefact masks) were cropped into 2048×2048-pixel image tiles. After class balancing, a total of 3,787 tiles were split 9:1 into training (3,409) and validation (378) sets. Tissue images differed with respect to the channels that were affected by artefacts (**Supplementary** Fig. 3a). The number of tiles containing artefacts also differed between images, ranging from as many as 463 tiles in image 59 to as few as 129 in image 64 (**Supplementary** Fig. 3b). Of the 3,787 total tiles, 1,734 contained pixels annotated as artefacts. Across all tiles, the average percentage of pixels affected by artefacts was ∼6.7% (**Supplementary** Fig. 1c).

Our DL model comprised a pretrained ResNet34 encoder^50^ coupled to a Feature Pyramid Network (FPN)^51^ decoder (ResNet-FPN). The input of the model were image tiles and its output was predicted binary artefact masks. To assess the technical reproducibility of artefact predictions, three independent ResNet-FPN models were trained to convergence starting from FPN network weights initialized using different random seeds. Validation loss (measured via Dice similarity coefficient) ranged from 0.426 to 0.459 (mean = 0.444). To determine the ability of the trained models to generalize across different marker channels, testing was performed on channel 29 of tissue section 54 (**Supplementary** Fig. 3d), which contained artefacts not found in other sections or channels (**Supplementary** Fig. 3a). Performance was assessed by precision-recall (PR) and receiver operating characteristic (ROC) curve analysis. Average precision (AP) ranged from 0.30 to 0.33 for the three models (**Supplementary** Fig. 3e) and area under the ROC curve (AUC) ranged between 0.71 and 0.75 (**Supplementary** Fig. 3f). This demonstrates that the assembly of a DL model for artefact detection in high-plex tissue images is feasible. However, we judge the overall level of performance relative to human reviewers to be inadequate and we strongly suspect that this is due to insufficient training data. CyLinter is nevertheless an ideal way to generate additional training data. Thus, we have established a deposition site at the Synapse data repository (Sage Bionetworks, https://www.synapse.org/#!Synapse:syn24193163/wiki/624232) for collecting CyLinter-curated image artefacts. We anticipate that further training of our ResNet-FPN model on this corpus of collected artefacts will ultimately yield a highly-performant model for integration into future iterations of the CyLinter workflow.

**Online Supplementary** Fig. 1 **Example artefacts in Dataset 1 (TOPACIO) (**https://www.synapse.org/#!Synapse:syn53781614**). a**, Twelve (12) examples of tissue folds. **b**, Twelve (12) examples of slide debris. **c**, Four (4) examples of coverslip air bubbles. **d**, Twelve (12) examples of image blur.

**Online Supplementary** Fig. 2 **Image galleries of clustering cells from pre-QC Dataset 2 (CRC) (**https://www.synapse.org/#!Synapse:syn53781627). Twenty (20) examples of cells from each of 22 clusters identified in the pre-QC CRC dataset showing the top three most highly expressed markers (1: green, 2: red, 3: blue) and Hoechst dye (gray). A single white pixel at the center of each image highlights the reference cell. Nuclear segmentation outlines are superimposed to show segmentation quality.

**Online Supplementary** Fig. 3 **Image galleries of clustering cells from pre-QC Dataset 6 (CODEX) (**https://www.synapse.org/#!Synapse:syn53781635). Twenty (20) examples of cells from each of 32 clusters identified in the pre-QC CODEX dataset (normal large intestine, specimen 1) showing the top three highly expressed markers (1: green, 2: red, 3: blue) and Hoechst dye (gray). A single white pixel at the center of each image highlights the reference cell. Nuclear segmentation outlines are superimposed to show segmentation quality.

**Online Supplementary** Fig. 4 **Image galleries of clustering cells from pre-QC Dataset 1 (TOPACIO) (**https://www.synapse.org/#!Synapse:syn53782191). Twenty (20) examples of cells from each of 48 (of 492) clusters identified in the pre-QC TOPACIO dataset showing the top three most highly expressed markers (1: green, 2: red, 3: blue) and Hoechst dye (gray). A single white pixel at the center of each image highlights the reference cell. Nuclear segmentation outlines are superimposed to show segmentation quality.

**Online Supplementary** Fig. 5 **Image tiles from Dataset 1 (TOPACIO) (**https://www.synapse.org/#!Synapse:syn53779745). Down-sampled, single-channel images from the 25 TNBC tissue specimens analyzed in this study used to estimate the number of image tiles impacted by microscopy artefacts. Artefact counts table and patient metadata table are also provided.

**Online Supplementary** Fig. 6 **Image galleries of clustered cells from post-QC Dataset 2 (CRC) (**https://www.synapse.org/#!Synapse:syn53781719). Twenty (20) examples of cells from each of 78 clusters identified in the post-QC CRC dataset showing the top three most highly expressed markers (1: green, 2: red, 3: blue) and Hoechst dye (gray). A single white pixel at the center of each image highlights the reference cell. Nuclear segmentation outlines are superimposed to show segmentation quality.

**Online Supplementary** Fig. 7 **Image galleries of clustered cells from post-QC Dataset 6 (CODEX) (**https://www.synapse.org/#!Synapse:syn53781730). Twenty (20) examples of cells from each of 28 clusters identified in the post-QC CODEX dataset showing the top three most highly expressed markers (1: green, 2: red, 3: blue) and Hoechst dye (gray). A single white pixel at the center of each image highlights the reference cell. Nuclear segmentation outlines are superimposed to show segmentation quality.

**Online Supplementary** Fig. 8 **Image galleries of clustered cells from post-QC Dataset 1 (TOPACIO) (**https://www.synapse.org/#!Synapse:syn53781892). Twenty (20) examples of cells from each of 43 clusters identified in the post-QC TOPACIO dataset showing the top three highly expressed markers (1: green, 2: red, 3: blue) and Hoechst dye (gray). A single white pixel at the center of each image highlights the reference cell. Nuclear segmentation outlines are superimposed to show segmentation quality.

**Online Supplementary** Fig. 9 **Channel colormaps applied to cells in the pre-QC Dataset 6 (CODEX) embedding (**https://www.synapse.org/#!Synapse:syn53781812).

**Online Supplementary** Fig. 10 **Channel colormaps applied to cells in the post-QC Dataset 6 (CODEX) embedding (**https://www.synapse.org/#!Synapse:syn53781819).

## METHODS

### Software Implementation

CyLinter software is written in Python3, archived on the Anaconda package repository, versioned controlled on Git/GitHub (https://github.com/labsyspharm/cylinter), instantiated as a configurable Python Class object, and validated for Mac, PC, and Linux operating systems. The tool can be installed at the command line using the Anaconda package installer (see the CyLinter website: https://labsyspharm.github.io/cylinter/ for details) and is executed with the following command: *cylinter configuration.yml,* where configuration.yml is an experiment-specific YAML configuration file. An optional *--module* flag can be passed before specifying the path to the configuration file to begin the pipeline at a specified module. More details on configuration settings can be found at the CyLinter website and GitHub repository (https://github.com/labsyspharm/cylinter^52^). The tool uses the Napari image viewer^53^ for image browsing and annotation tasks. The tool also uses numerical and image-processing routines from multiple Python data science libraries, including pandas, numpy, matplotlib, seaborn, SciPy, scikit-learn, and scikit-image. OME-TIFF files are read using tifffile and processed into multi-resolution pyramids using a combination of Zarr and dask routines that allow for rapid panning and zooming of large (hundreds of GB) images. The CyLinter pipeline consists of multiple QC modules, each implemented as a Python function, that perform different visualization, data filtration, or analysis tasks. Several modules return redacted versions of the input spatial feature table, while others perform analysis tasks such as cell clustering.

CyLinter is freely-available for academic re-use under the MIT license. A minimal example dataset consisting of 4 tissue cores from the EMIT TMA22 dataset can be downloaded from the Synapse data repository (Synapse ID: syn52468155) by following instructions at the CyLinter website (https://labsyspharm.github.io/cylinter/exemplar/). All CyLinter analyses presented in this work were performed on a commercially available 2019 MacBook Pro equipped with eight 2.4 GHz Intel Core i9 processors (5.0GHz Turbo Boost) and 32GB 2400MHz DDR4 memory. Imaging data analyzed in this study were stored on and accessed from an external hard drive with 12TB capacity. Implemented software versions are as follows: Python 3.11.5, CyLinter 0.0.47.

### t-CyCIF

The CyCIF approach to multiplex imaging involves iterative cycles of antibody incubation with tissue, imaging, and fluorophore deactivation as described previously^24^; protocols and methods related to CyCIF are available on Protocols.io (see “Detailed Experimental Protocols” below). Briefly, multiplex CyCIF images were collected using a RareCyte CyteFinder II HT Instrument equipped with a 20x (0.75 NA) objective and 2×2 pixel binning. This setup allowed for the acquisition of 4-channel image tiles with dimensions 1280×1080 pixels and a corresponding pixel size of 0.65 μm/pixel. All four channels are imaged during each round of CyCIF, one of which is always reserved for nuclear counterstain (Hoechst or DAPI) to visualize cell nuclei. RCPNL files containing 16-bit imaging data were generated (one per image tile) during each imaging cycle.

### Image Processing

Raw microscopy image tiles (RCPNL files) for the datasets described in this study were processed into stitched, registered, and segmented OME-TIFF^54^ files using the MCMICRO image-processing pipeline^28^. Corresponding cell x feature CSV files (i.e., spatial feature tables) were also generated by MCMICRO. Specific algorithms implemented in MCMICRO for the processing of each dataset are as follows: BaSiC—a Fiji/ImageJ plugin for background and shading correction used to perform flatfield and darkfield image correction^55^; ASHLAR—a program for seamless mosaic image processing across imaging cycles^37^; Coreograph (used for the EMIT dataset, https://github.com/HMS-IDAC/UNetCoreograph)—for dearraying the mosaic TMA image into individual TIFF and CSV files per core; UnMICST^38^—used for cell segmentation; employs the U-Net^56^ deep learning architecture for semantic segmentation; S3segmenter (https://github.com/HMS-IDAC/S3segmenter); MCQuant (https://github.com/labsyspharm/quantification) for per cell feature extraction including X,Y spatial coordinates, segmentation areas, mean marker intensities, and nuclear morphology attributes.

### Automated Artefact Detection in CyLinter with Classical Algorithms

An algorithm consisting of classical image analysis steps was designed to automatically identify prevalent artefacts commonly found in highly multiplexed images (e.g., illumination aberrations, antibody aggregates, and tissue folding). The model is applied on a channel-by-channel basis and works on down-sampled versions of each channel, rescaling pixel values to uint8 bit depth for efficient processing. A series of operations in mathematical morphology consisting of erosion and local mean smoothing followed by dilation are applied to transform each down-sampled image channel. These three steps utilize a disk kernel, where the kernel size is a user-defined parameter assumed to have a diameter on the order of 3-5 single cells, conditional on image pixel size. This kernel is then expanded to find local maxima seed points corresponding to putative artefacts. Each artefact is extracted via a flood fill operation according to a specific tolerance parameter that is adjusted in real-time by the user. The union of the flood fill regions produces a binary artefact mask that is resized to the original image dimensions; cells falling within mask boundaries are then dropped from the corresponding spatial feature table.

### Deep Learning-based Automated Artefact Detection

The machine learning artefact detection model implemented in this study derives from the Feature Pyramid Network (FPN)^51^, a fully convolutional encoder-decoder architecture designed for object detection tasks applicable to semantic image segmentation. The encoder network is implemented using a ResNet34 backbone^50^ with model parameters initialized from the pretraining weights on ImageNet. Input image tiles of size 2048×2048-pixels (acquired at a nominal resolution of 0.65µm/pixel) were down-sampled to 256×256-pixels and fed into the encoder network to produce low-resolution feature maps. Resulting feature maps were then decoded into feature pyramids through iterated up-sampling using a bilinear interpolation and combined with the original feature maps. Each layer of the feature pyramid was up-sampled to the same resolution and segmented such that all resulting predicted artefact masks were combined to yield the final composite prediction mask. The FPN architecture is implemented using the Segmentation Models library for image segmentation based on the Python and PyTorch frameworks^57^. The model was trained using the Adam optimizer with a DICE loss function and a fixed learning rate (1×10^-4^) using a batch size of 16 image tiles for 10 epochs.

### Dataset 1 (TOPACIO, CyCIF)

The TOPACIO dataset used in this study consists of 25 de-identified formalin-fixed, paraffin embedded (FFPE) tissue sections (5 μm thick) of triple-negative breast cancer from patients enrolled in the TOPACIO clinical trial (ClinicalTrials.gov Identifier: NCT02657889). Specimens were collected via one of three different biopsy methods: fine needle, punch needle, or gross tumor resection and procured from Tesaro and Merck & Co., Inc., Rahway, NJ, USA as part of the recently-completed trial. Slides were mounted onto Superfrost Plus glass microscope slides (Fisher Scientific, 12-550-15) then dewaxed and antigen-retrieved using a Leica BOND RX Fully Automated Research Stainer prior to multiplex data acquisition by CyCIF. Images were acquired at 20x magnification with 2×2 binning (0.65 μm/pixel nominal resolution) over 10 CyCIF cycles using 27 markers (19 plus Hoechst evaluated in this study); see **Supplementary Table 1** for further details. Clustering of cells in this dataset (totaling ∼1.9×10^7^ segmented nuclei) was performed on a randomly selected subset of ∼3×10^6^ cells to reduce computing time.

### Dataset 2 (CRC, CyCIF)

The CRC dataset consists of a whole-slide section (1.6cm^2^) of human colorectal adenocarcinoma tissue (section# 097) from a 69-year-old white male imaged at 20x magnification with 2×2 binning (0.65 μm/pixel nominal resolution) over 10 CyCIF cycles using 24 markers across 10 CyCIF cycles (21 plus Hoechst evaluated in the current study) collected as part of the Human Tumor Atlas Network (HTAN); see **Supplementary Table 1** for further details.

### Dataset 3 (EMIT TMA22, CyCIF)

The EMIT TMA dataset consists of human tissue specimens from 42 patients organized as a multi-tissue microarray (HTMA427) under an excess tissue protocol (clinical discards) approved by the IRB at Brigham and Women’s Hospital (BWH IRB 2018P001627). Two (2) 1.5 mm diameter cores were acquired from each of 60 tissue regions with the goal of acquiring one or two examples of as many tumors as possible (with matched normal tissue from the same resection when feasible). Overall, the TMA contains 123 cores including 3 “marker cores” consisting of normal kidney cortex which were added to the TMA in an arrangement that makes it possible to orient the overall TMA image. Not including the marker cores 44 cores were from males and 76 were from females between 21 and 86 years-of-age. The EMIT TMA22 dataset was acquired at 20x magnification with 2×2 binning (0.65 μm/pixel nominal resolution) over 10 CyCIF cycles using 27 markers (20 plus Hoechst evaluated in the current study) and is available for download from the Synapse data repository (https://www.synapse.org/#!Synapse:syn22345750); see **Supplementary Table 1** for further details.

### Dataset 4 (HNSCC, CODEX)

The HNSCC CODEX dataset consists of two sections of the same deidentified specimen of head & neck squamous carcinoma (HNSCC) imaged at 20x magnification with 2×2 binning (0.65 μm/pixel nominal resolution) over 9 imaging cycles using 15 markers plus DAPI; see **Supplementary Table 1** for further details.

### Dataset 5 (normal tonsil, mIHC)

The mIHC dataset consists of a deidentified whole-slide tonsil specimen from a 4-year-old female of European ancestry procured from the Cooperative Human Tissue Network (CHTN), Western Division, as part of the HTAN SARDANA Trans-Network Project and imaged at 20x magnification with 2×2 binning (0.5 μm/pixel nominal resolution) over 5 mIHC cycles using 18 markers plus Hoechst; see **Supplementary Table 1** for further details.

### Dataset 6 (normal large intestine, CODEX, specimen 1)

A single section of deidentified human tissue from a 78-year-old African American male imaged at 20x magnification (0.75NA, 0.38 μm/pixel nominal resolution) over 23 imaging cycles using 59 markers (58 evaluated in this study, as DRAQ5 was excluded due to its overlap with Hoechst). These data were collected at Stanford University as part of the Human BioMolecular Atlas Program (HuBMAP); see **Supplementary Table 1** for further details.

### Dataset 7 (normal large intestine, CODEX, specimen 2)

The large intestine CODEX dataset consists of a single section of deidentified human tissue from a 24-year-old white male imaged at 20x magnification (0.75NA, 0.38 μm/pixel nominal resolution) over 24 imaging cycles using 54 markers (53 evaluated in this study, as DRAQ5 was excluded due to its overlap with Hoechst). These data were collected at Stanford University as part of the Human BioMolecular Atlas Program (HuBMAP); see **Supplementary Table 1** for further details.

### Detailed Experimental Protocols

1. FFPE Tissue Pre-treatmet Before t-CyCIF on Leica Bond RX V.2 (dx.doi.org/10.17504/protocols.io.bji2kkge)
2. Tissue Cyclic Immunofluorescence (t-CyCIF) V.2 (dx.doi.org/10.17504/protocols.io.bjiukkew)

### Ethics and IRB Statement

Tissue specimens from the recently completed TOPACIO clinical trial (ClinicalTrials.gov Identifier: NCT02657889) which was conducted in accordance with ethical principles founded in the Declaration of Helsinki. This study received central approval by the Dana-Farber institutional review board, and/or relevant competent authorities at each site. All patients provided written informed consent to participate in the study. All specimens and data have been deidentified for the work performed at Harvard Medical School, approved under Institutional Review Boards (IRB) protocol 19-0186. The research complies with all relevant ethical regulations, was reviewed and approved by the IRBs at HMS and DFCI and is considered Non-Human Subjects Research.

### Reporting Summary

Further information on research design is available in the Nature Portfolio Reporting Summary linked to this article.

### Data Availability Statement

New data associated with this paper is available at the HTAN Data Portal (https://data.humantumoratlas.org). Previously published data is through public repositories. See **Supplementary Table 1** for a complete list of datasets and their associated identifiers and repositories. **Online Supplementary** Figures 1-4 and the CyLinter demonstration dataset can be accessed at Sage Synapse (https://www.synapse.org/#!Synapse:syn24193163/files)

### Code Availability Statement

CyLinter source code is available for academic re-use under the MIT open-source license agreement at Github (https://github.com/labsyspharm/cylinter)^52^. Python code used to generate the findings of the study is available at https://github.com/labsyspharm/cylinter-paper which is archived on Zenodo at https://zenodo.org/doi/10.5281/zenodo.10067803.

### Author Contribution Statements

G.J.B conceived and designed the study. P.K.S. supervised the work and secured funding. G.J.B. developed the CyLinter software, J.L.G and E.A.M. provided access to the TOPACIO clinical biopsies, J.R.L acquired t-CyCIF data from the TOPACIO specimens, T.V. and J.D. curated tissue ROIs for the TOPACIO specimens, E.N. and Z.Z. developed the method for automated artefact detection, G.J.B performed CyLinter analysis on all datasets and generated the figures, G.J.B and P.K.S wrote the manuscript with input from all authors.

## Acknowledgements

This work was supported by the Ludwig Cancer Research and the Ludwig Center at Harvard (P.K.S., S.S.) and by NIH NCI grants U2C-CA233280, and U2C-CA233262 (P.K.S., S.S.). Development of computational methods and image processing software is supported by a Team Science Grant from the Gray Foundation (P.K.S., S.S.), the Gates Foundation grant INV-027106 (P.K.S.), the David Liposarcoma Research Initiative at Dana-Farber Cancer Institute supported by KBF Canada via the Rossy Foundation Fund (P.K.S., S.S.) and the Emerson Collective (P.K.S.). S.S. is supported by the BWH President’s Scholars Award. We gratefully acknowledge Juliann Tefft for superb editorial support; Clarence Yapp for help with artifact annotation; Kai Wucherpfennig and Sascha Marx for providing the HNSCC CODEX dataset; Zoltan Maliga and Connor Jacobson for providing CyCIF EMIT TMA22 images; and the Dana-Farber/Harvard Cancer Center for use of the Specialized Histopathology Core, which provided TMA construction and sectioning services. We also thank Yu-An Chen for assisting in the collection of CyCIF data from the SARDANA-097 tissue specimen performed as part of the NCI Human Tumor Atlas Network (HTAN).

## Competing Interests

P.K.S. is a cofounder and member of the Board of Directors of Glencoe Software, a member of the Board of Directors for Applied Biomath and a member of the Scientific Advisory Board for RareCyte, NanoString and Montai Health; he holds equity in Glencoe and RareCyte. P.K.S. is a consultant for Merck. PKS declares that none of these relationships have influenced the content of this manuscript. E. A. M. reports compensated service on Scientific Advisory Boards for Astra Zeneca, BioNTech and Merck; uncompensated service on Steering Committees for Bristol Myers Squibb and Roche/Genentech; speakers’ honoraria and travel support from Merck Sharp & Dohme; and institutional research support from Roche/Genentech (via an SU2C grant) and Gilead. She also reports research funding from Susan Komen for the Cure for which she serves as a Scientific Advisor, and uncompensated participation as a member of the American Society of Clinical Oncology Board of Directors. J. L. G. serves or has previously served on advisory boards and/or as a scientific advisory board member for Array BioPharma/Pfizer, AstraZeneca, BD Biosciences, Carisma, Codagenix, Duke Street Bio, GlaxoSmithKline, Kowa, Kymera, OncoOne and Verseau Therapeutics, and has research grants from Array BioPharma/Pfizer, Duke Street Bio, Eli Lilly, GlaxoSmithKline and Merck. The other authors declare no competing interests.

---

**Supplementary Fig. 1.**
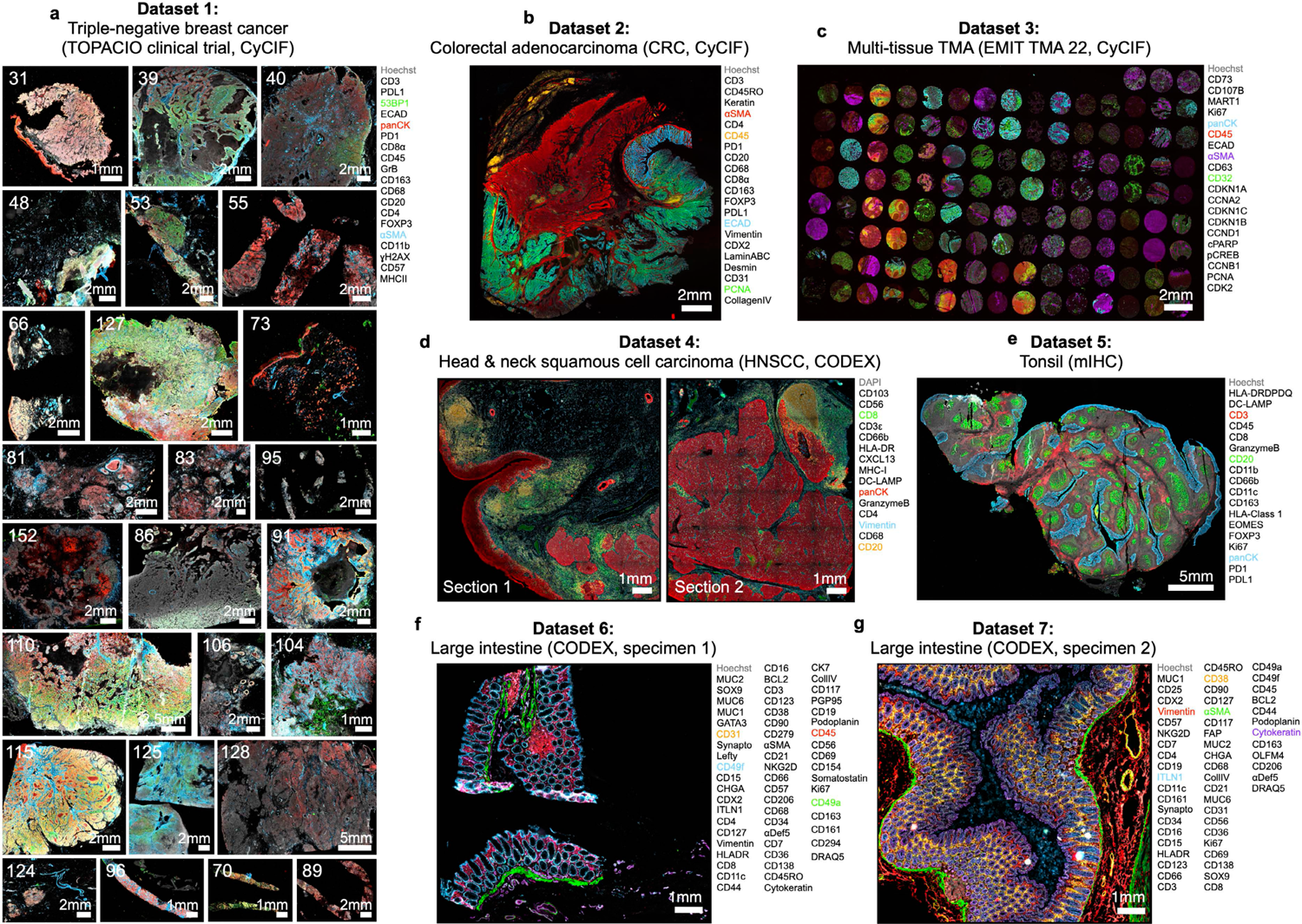
Overview of the seven multiplex IF datasets analyzed in this study. **a**, Dataset 1 (TOPACIO, CyCIF): 25 human TNBC clinical trial specimens (∼6-353 mm^2^). Numbers in upper left of each panel indicate specimen number. Channels shown are Hoechst (gray), 53BP1 (green), panCK (red), and αSMA (blue). **b**, Dataset 2 (CRC, CyCIF): an ∼172 mm^2^ whole-slide section of primary human colorectal adenocarcinoma. Channels shown are Hoechst (gray), αSMA (red), CD45 (orange), ECAD (blue), and PCNA (green). **c**, Dataset 3 (EMIT TMA22, CyCIF): 123 healthy and diseased human tissue cores each ∼2 mm^2^ arranged on a single microscope slide. Channels shown are Hoechst (gray), panCK (blue), CD45 (red), αSMA (purple), and CD32 (green). **d**, Dataset 4 (HNSCC, CODEX): two ∼42 mm^2^ whole-slide sections of human HNSCC. Channels shown are DAPI (gray), CD8 (green), panCK (red), vimentin (blue), and CD20 (orange). **e**, Dataset 5 (Tonsil, mIHC): an ∼92 mm^2^ whole-slide section of normal human tonsil. Channels shown are Hoechst (gray), CD3 (red), CD20 (green), panCK (blue). **f**, Dataset 6 (Large intestine, CODEX, specimen 1): an ∼7 mm^2^ whole-slide section of normal human large intestine from a 78-year-old African American male. Channels shown are Hoechst (gray), CD31 (orange), CD49f (blue), CD45 (red), CD49a (green). **g**, Dataset 7 (Large intestine, CODEX, specimen 2): an ∼12 mm^2^ whole-slide section of normal human large intestine from a 24-year-old white male. Channels shown are Hoechst (gray), Vimentin (red), ITLN1 (blue), CD38 (orange), αSMA (green), Cytokeratin (purple). Markers to the right of each dataset indicate the full marker set captured in the corresponding image(s). See **Supplementary Table 1** for specimen identifiers and data access information.

**Extended Data Fig. 1.**
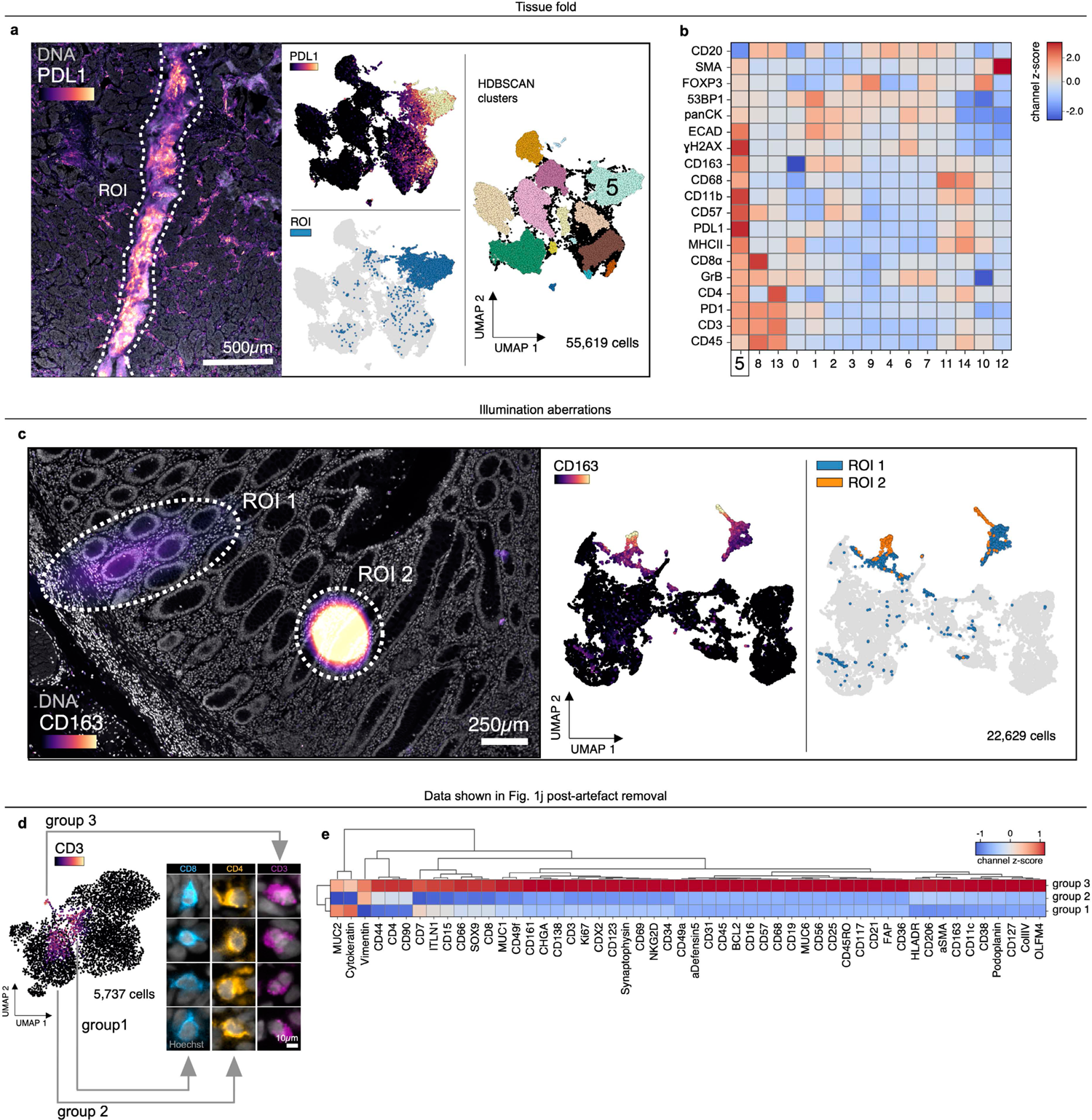
Recurring artefacts in whole slide immunofluorescence images of tissue and their effects on tissue-derived single-cell data. **a**, Left: Field of view from Dataset 1 (TOPACIO, specimen 110) showing a tissue fold (ROI, dashed white outline) as viewed in channels PDL1 (colormap) and Hoechst (gray). Right: UMAP embedding of 19-channel single-cell data from the image at left colored by PDL1 intensity (top left), cells contained within the ROI (bottom left), and HDBSCAN cluster (center right). Cells in cluster 5 (labeled) are those affected by the tissue fold and form of a discrete cluster in UMAP space. **b**, Clustered heatmap showing channel z-scores for HDBSCAN clusters from panel (a) demonstrating that cluster 5 cells (those affected by the tissue fold) are artificially bright for all channels presumably due to a combination of tissue overlap and insufficient antibody washing. **c**, Left: Field of view from Dataset 2 (CRC) showing two illumination aberrations (ROIs, dashed white outlines) as viewed in channels CD163 (colormap) and Hoechst (gray). Right: UMAP embedding of 21-channel single-cell data from the image at left colored by CD163 intensity (left) and whether the cells fall within one of the two ROIs (right). **d**, UMAP embedding of the 52-channel single-cell data shown in **Fig. 1j** (Dataset 7, large intestine, CODEX) after cells affected by the five illumination aberrations have been removed. Three groups of cells bright for CD3 remain (groups 1-3). Image galleries at right show 4 examples of each cell type in representative channels: group 1 = CD8^+^ T cells, group 2 = CD4^+^ T cells, group 3 = undefined cells immunoreactive to all 52 channels (not due to microscopy artefacts). **e**, Clustered heatmap showing channel z-scores for HDBSCAN clusters from panel (d) demonstrating that group 3 cells are bright for all 52 channels despite not being affected by microscopy artefacts.

**Extended Data Fig. 2.**
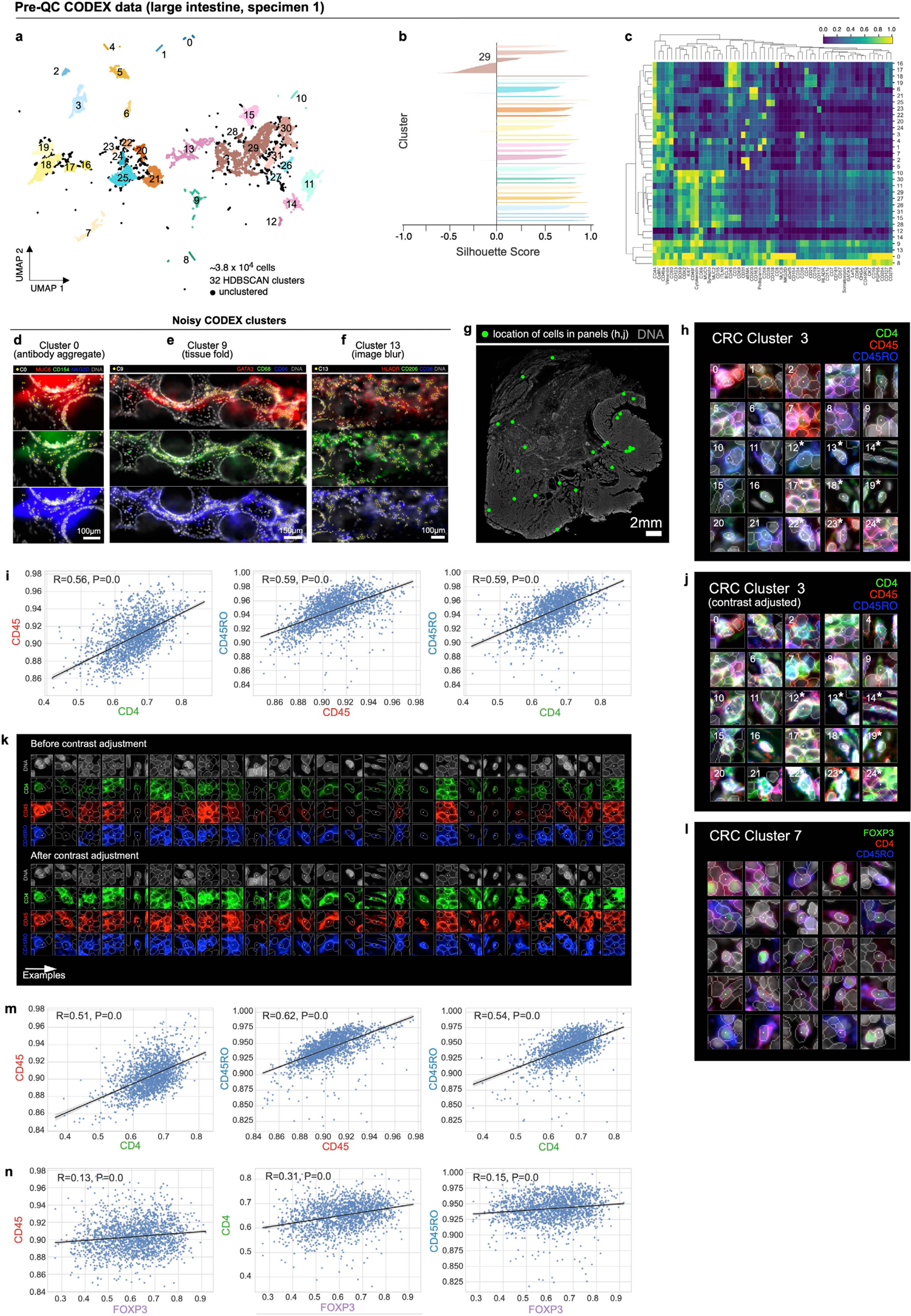
Evaluation of pre-QC cell clustering results from Dataset 6 (large intestine, CODEX) and Dataset 2 (CRC, CyCIF). **a**, UMAP embedding of Dataset 6 showing ∼3.8×10^4^ cells colored by HDBSCAN cluster (numbered 0-31). Black scatter points represent unclustered cells (10.5% of cells). **b**, Silhouette scores for CODEX clusters shown in panel (a). Cluster 29 exhibits cells with negative silhouette scores indicative of under-clustering. **c**, Clustered heatmap of clusters from Dataset 6 showing mean signal intensities of clustering cells normalized across clusters (row-wise). **d**, Correlated, non-specific signals in a region of Dataset 6 as seen in channels MUC6 (red), CD154 (green), and NKG2D (blue). Yellow dots highlight cluster 0 cells which have formed due to this artefact; Hoechst (gray) shown for reference. **e**, Tissue fold in a region of Dataset 6 as seen in channels GATA3 (red), CD68 (green), and CD66 (blue). Yellow dots highlight cluster 9 cells which have formed due to this artefact; Hoechst (gray) shown for reference. **f**, Image blur in a region of Dataset 6 as seen in channels HLADR (red), CD206 (green), and CD38 (blue). Yellow dots highlight cluster 13 cells which have formed due to this artefact; Hoechst (DNA, gray) shown for reference. **g**, Location of CRC cluster 3 cells shown in panel (g) revealing no regional bias in the distribution of cells. **h**, Top three most highly expressed markers (1: green, 2: red, 3: blue) for the 25 members of CRC cluster 3 (memory helper T) cells represented by the rugplots of **Fig. 2n**. White asterisks highlight cells shown in enlarged format in **Fig. 2m**. A single white pixel at the center of each image patch highlights the reference cell. Nuclear segmentation outlines (translucent white outlines) and Hoechst (gray) shown for reference. **i**, Regression plots showing correlation (two-sided, Pearson R, p < 0.05) among CD4, CD45, and CD45RO marker expression by 1.9×10^3^ CRC cluster 3 cells. **j**, CRC cluster 3 cells shown in panel (h) after signal intensity cutoffs have been adjusted per image to improve the homogeneity of their appearance. White asterisks highlight cells shown in enlarged format in (**Fig. 2o**). **k**, CRC cluster 3 cells shown in panels (h) and (j) with channels shown separately for clarity: Hoechst (gray), CD4 (green), CD45 (red), CD45RO (blue). Top panels show cells before contrast adjustment (panel h), bottom panels show cells after contrast adjustment (panel j). **l**, Top three most highly expressed markers (1: green, 2: red, 3: blue) for 25 CRC cluster 7 (Treg) cells. A single white pixel at the center of each image patch highlights the reference cell. Nuclear segmentation outlines (translucent white outlines); Hoechst (gray) shown for reference. **m**, Regression plots showing strong correlation (two-sided, Pearson R, p < 0.05) among CD4, CD45, and CD45RO marker expression of 1.9×10^3^ CRC cluster 7 cells. **n**, Regression plots showing weak correlation (two-sided, Pearson R, p < 0.05) between FOXP3 and CD4, CD45, and CD45RO marker expression of 1.9×10^3^ CRC cluster 7 cells.

**Extended Data Fig. 3.**
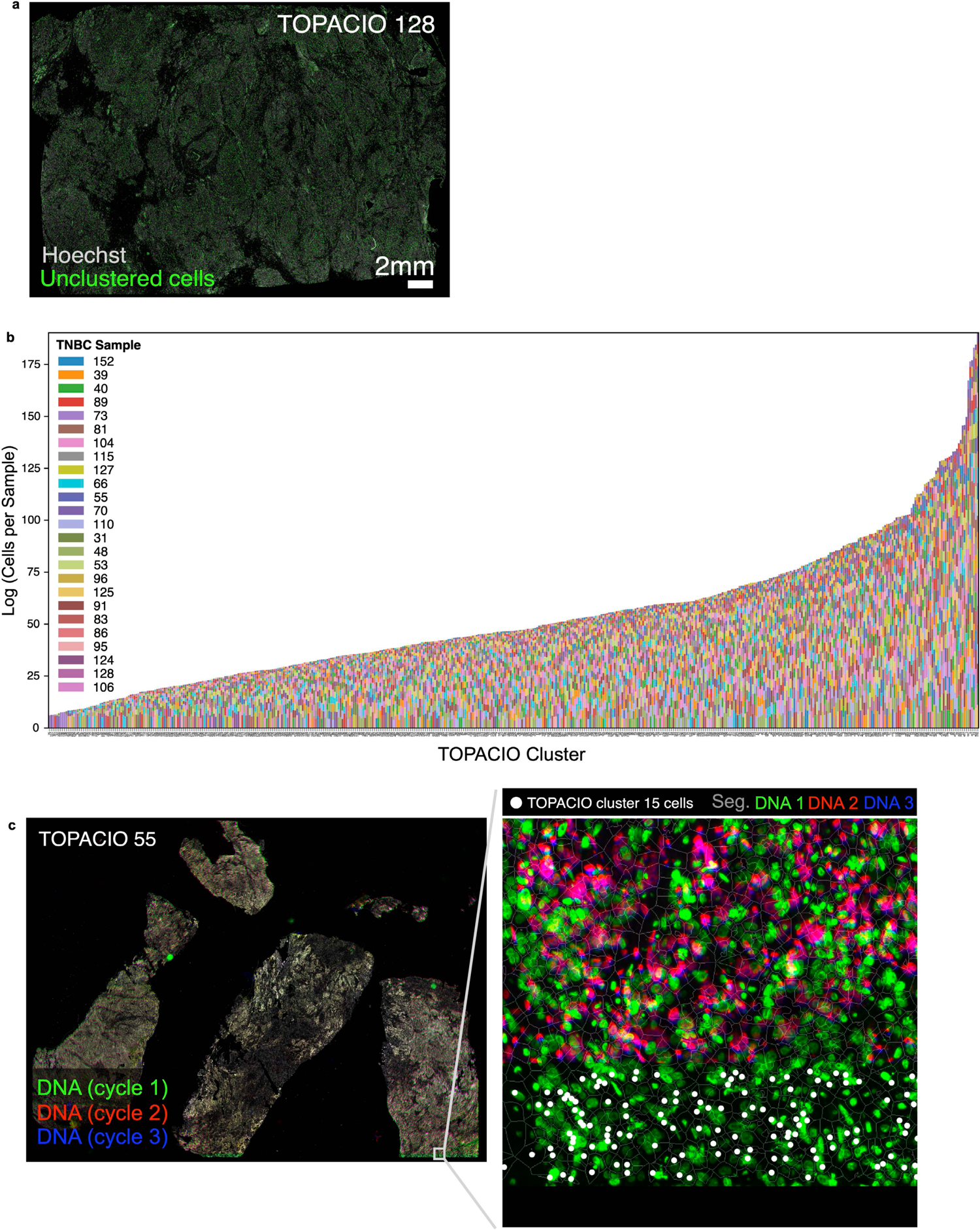
Evaluation of pre-QC cell clustering results from Dataset 1 (TOPACIO). **a**, Spatial distribution of unclustered (ambiguous) cells (green dots) from the pre-QC TOPACIO embedding shown in **Fig. 3a** as represented by specimen 55, which exhibits no discernable spatial pattern of sampling bias; Hoechst (gray) shown for reference. **b**, Stacked bar charts showing the relative contribution of each patient specimen to each cluster. **c**, TOPACIO specimen 55 at low (left) and high (right) magnification showing Hoechst signals for the first three imaging cycles: cycles 1 (green), 2 (red), and 3 (blue) have been superimposed to demonstrate a cross-cycle image alignment problem at the bottom of this specimen. Small white box at the bottom-right of the low magnification image shows the location of the higher magnification image. White dots in the high magnification image highlight TOPACIO cluster 15 cells which have formed due to this image alignment artefact.

**Extended Data Fig. 4.**
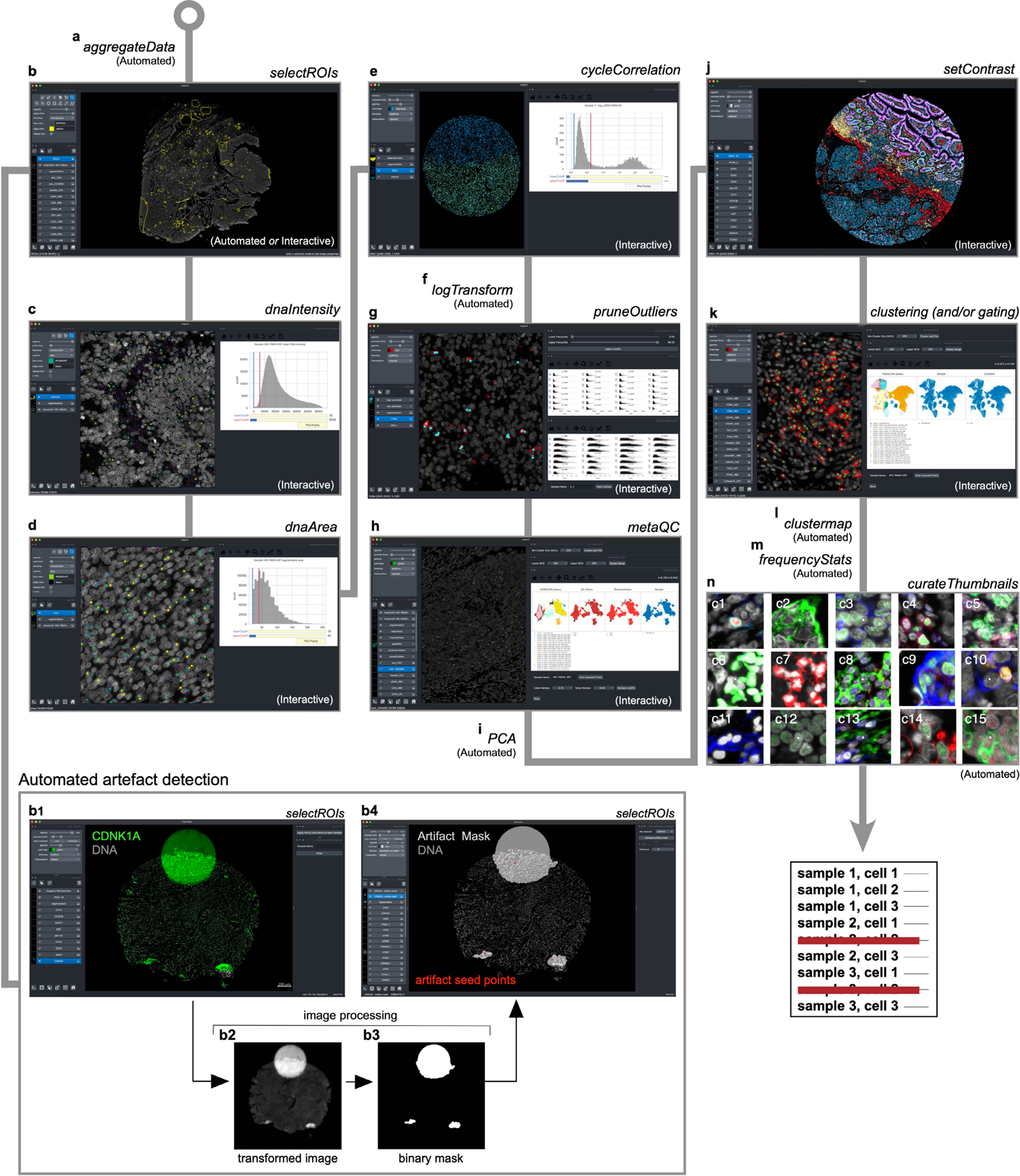
Identifying and removing noisy single-cell data points with CyLinter. CyLinter workflow (see project website for implementation details: https://labsyspharm.github.io/cylinter/modules/). **a**, Aggregate data (automated): raw spatial feature tables for all specimens in a batch are merged into a single Pandas (Python) dataframe. **b**, ROI selection (interactive *or* automated): multi-channel images are viewed to identify and gate on regions of tissue affected by microscopy artefacts (negative selection mode) or areas of tissue devoid of artefacts (positive selection mode. **b_1_-b_4_**, Demonstration of automated artefact detection in CyLinter: **b_1_**, CyLinter’s *selectROIs* module showing artefacts in the CDKN1A (green) channel of Dataset 3 (EMIT TMA, core 18, mesothelioma). **b_2_**, Transformed version of the original CDKN1A image such that artefacts appear as large, bright regions relative to channel intensity variations associated with true signal of immunoreactive cells which are suppressed. **b_3_**, Local intensity maxima are identified in the transformed image and a flood fill algorithm is used to create a pixel-level binary mask indicating regions of tissue affected by artefacts. In this example, the method identifies three artefacts in the image: one fluorescence aberration at the top of the core, and two tissue folds at the bottom of the core. **b_4_**, CyLinter’s *selectROIs* module showing the binary artefact mask (translucent gray shapes) and their corresponding local maxima (red dots) defining each of the three artefacts. **c**, DNA intensity filter (interactive): histogram sliders are used to define lower and upper bounds on nuclear counterstain single intensity. Cells between cutoffs are visualized as scatter points at their spatial coordinates in the corresponding tissue for gate confirmation or refinement. **d**, Segmentation area filter (interactive): histogram sliders are used to define lower and upper bounds on cell segmentation area (pixel counts). Cells between cutoffs are visualized as scatter points at their spatial coordinates in the corresponding tissue for gate confirmation or refinement. **e**, Cross-cycle correlation filter (interactive): applicable to multi-cycle experiments. Histogram sliders are used to define lower and upper bounds on the log-transformed ratio of DNA signals between the first and last imaging cycles. Cells between cutoffs are visualized as scatter points at their spatial coordinates in their corresponding tissues for gate confirmation or refinement. **f**, Log transformation (automated): single-cell data are log-transformed. **g**, Channel outliers filter (interactive): the distribution of cells according to antibody signal intensity is viewed for all specimens as a facet grid of scatter plots (or hexbin plots) against cell area (y-axes). Lower and upper percentile cutoffs are applied to remove outliers. Outliers are visualized as scatter points at their spatial coordinates in their corresponding tissues for gate confirmation or refinement. **h**, MetaQC (interactive): unsupervised clustering methods (UMAP or TSNE followed by HDBSCAN clustering) are used to correct for gating bias in prior data filtration modules by thresholding on the percent of each cluster composed of clean (maintained) or noisy (redacted) cells. **i**, Principal component analysis (PCA, automated): PCA is performed and Horn’s parallel analysis is used to determine the number of PCs associated with non-random variation in the dataset. **j**, Image contrast adjustment (interactive): channel contrast settings are optimized for visualization on reference tissues which are applied to all specimens in the cohort. **k**, Unsupervised clustering (interactive): UMAP (or TSNE) and HDBSCAN are used to identify unique cell states in a given cohort of tissues. Manual gating can also be performed to identify cell populations. **l**, Compute clustered heatmap (automated): clustered heatmap is generated showing channel z-scores for identified clusters (or gated populations). **m**, Compute frequency statistics (automated): pairwise t statistics on the frequency of each identified cluster or gated cell population between groups of tissues specified in CyLinter’s configuration file (*cylinter_config.yml*, e.g., treated vs. untreated, response vs. no response, etc.) are computed. **n**, Evaluate cluster membership (automated): cluster quality is checked by visualizing galleries of example cells drawn at random from each cluster identified in the *clustering* module (panel k).

**Extended Data Fig. 5.**
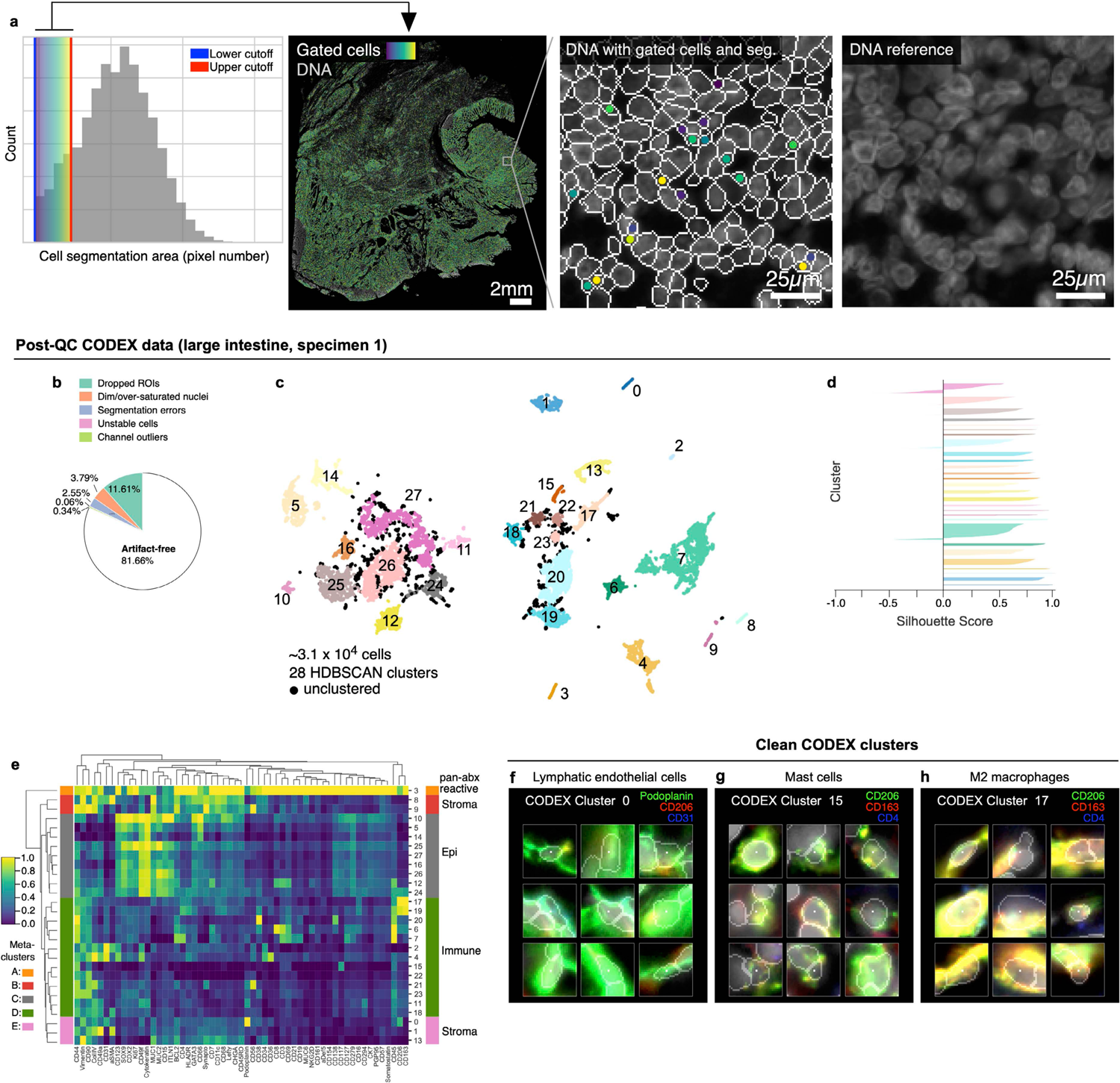
Over-segmentation in Dataset 2 (CRC, CyCIF) and Cleaning of Dataset 6 (large intestine, CODEX) with CyLinter. **a**, Gating of cells in the CRC image (Dataset 2) image according to nuclear segmentation area shows that this image contains several over-segmented nuclei (i.e., nuclei split into multiple segmentation objects). **b**, Fraction of cells in Dataset 6 (large intestine, CODEX, specimen 1) redacted by each QC filter in the CyLinter pipeline. Dropped ROIs = cells dropped by *selectROIs* module), Dim/over-saturated nuclei = cells dropped by *dnaIntensity* module), Segmentation errors = cells dropped by *areaFilter* module, Unstable cells = cells dropped by *cycleCorrelation* module, Channel outliers = cells dropped by *pruneOutliers* module, Artefact-free = cells remaining after QC. **c**, UMAP embedding of post-QC CODEX clusters showing ∼3.1×10^4^ cells colored by HDBSCAN cluster. Black scatter points represent unclustered cells (10.1% of cells). **d**, Silhouette scores for post-QC CODEX clusters shown in panel (c). **e**, Post-QC CODEX clustered heatmap showing mean signal intensities of clustering cells normalized across clusters (row-wise). Five (5) meta-clusters defined by the clustered heatmap dendrogram at the left are highlighted. **f-h**, Top three most highly expressed markers (1: green, 2: red, 3: blue) for clusters 0 (lymphatic endothelial cells, **f**), 15 (mast cells, **g**), and 17 (M2 macrophages, **h**). A single white pixel at the center of each image highlights the reference cell. Nuclear segmentation outlines (translucent outlines) and Hoechst (gray) shown for reference.

**Extended data Fig. 6.**
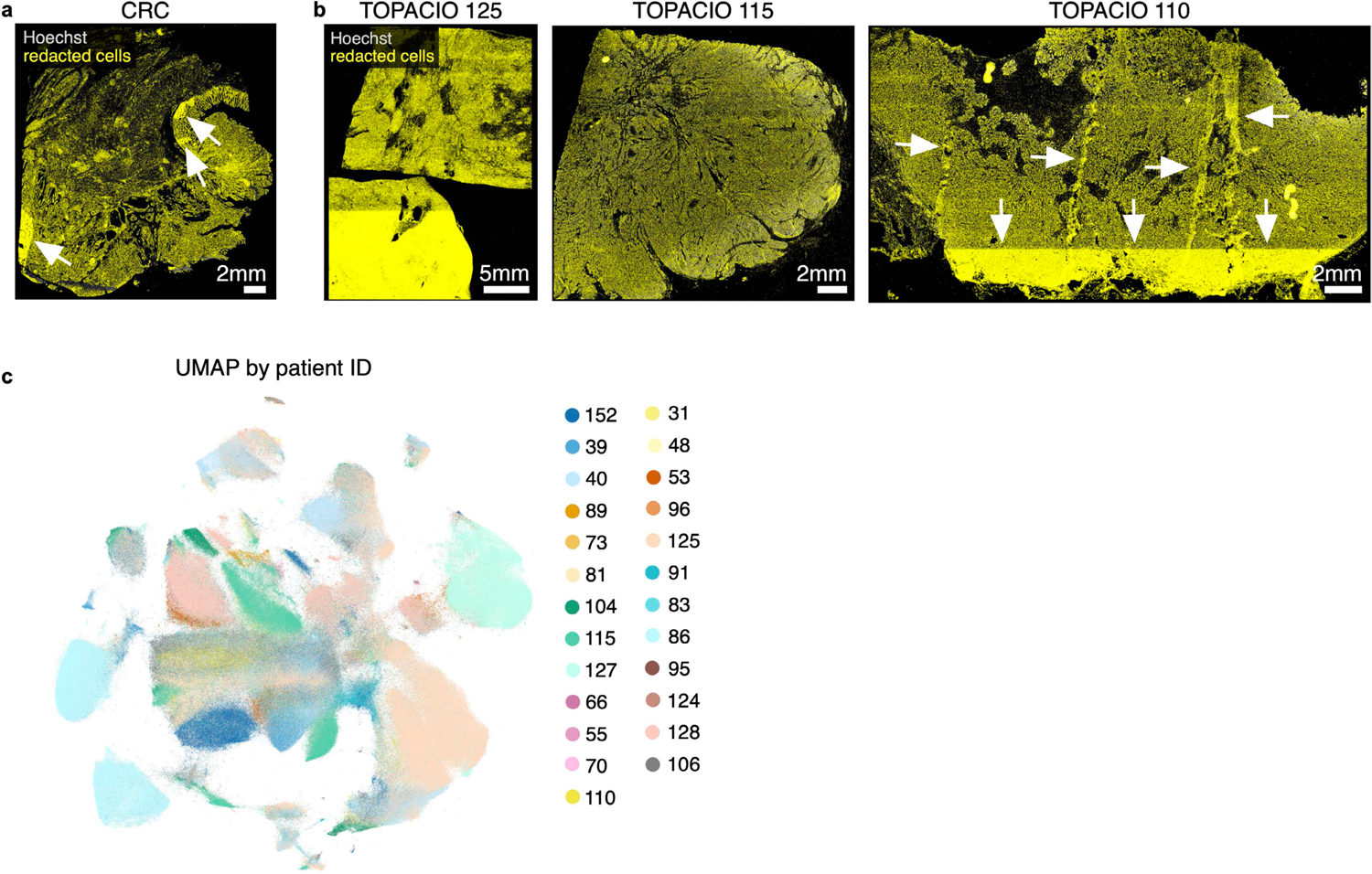
Location of cells redacted by CyLinter in Dataset 2 (CRC) and Dataset 1 (TOPACIO) and Post-QC TOPACIO UMAP embedding colored by patient ID. **a**, Cells redacted by CyLinter from the Dataset 2 (CRC) demonstrating no discernable bias in the removal of cells from the image with the exception of those areas highlighted by the white arrows which were affected by focal artefacts and removed using CyLinter’s *selectROIs* module. b, Cells redacted by CyLinter from three arbitrary specimens from Dataset 1 (TOPACIO) demonstrating no discernable bias in the removal of cells from the images with the exception of those areas highlighted by the white arrows which were affected by focal artefacts and removed using CyLinter’s *selectROIs* module. c, UMAP embedding of post-QC TOPACIO data shown in (**Fig. 6b**) colored by specimen ID demonstrating patient-specific clustering in tumor cell populations, but not immune and stromal populations (refer to **Fig. 6b,d**,e-i and Online Supplementary Fig. 8 for cluster phenotype identities).

**Supplementary Fig. 2.**
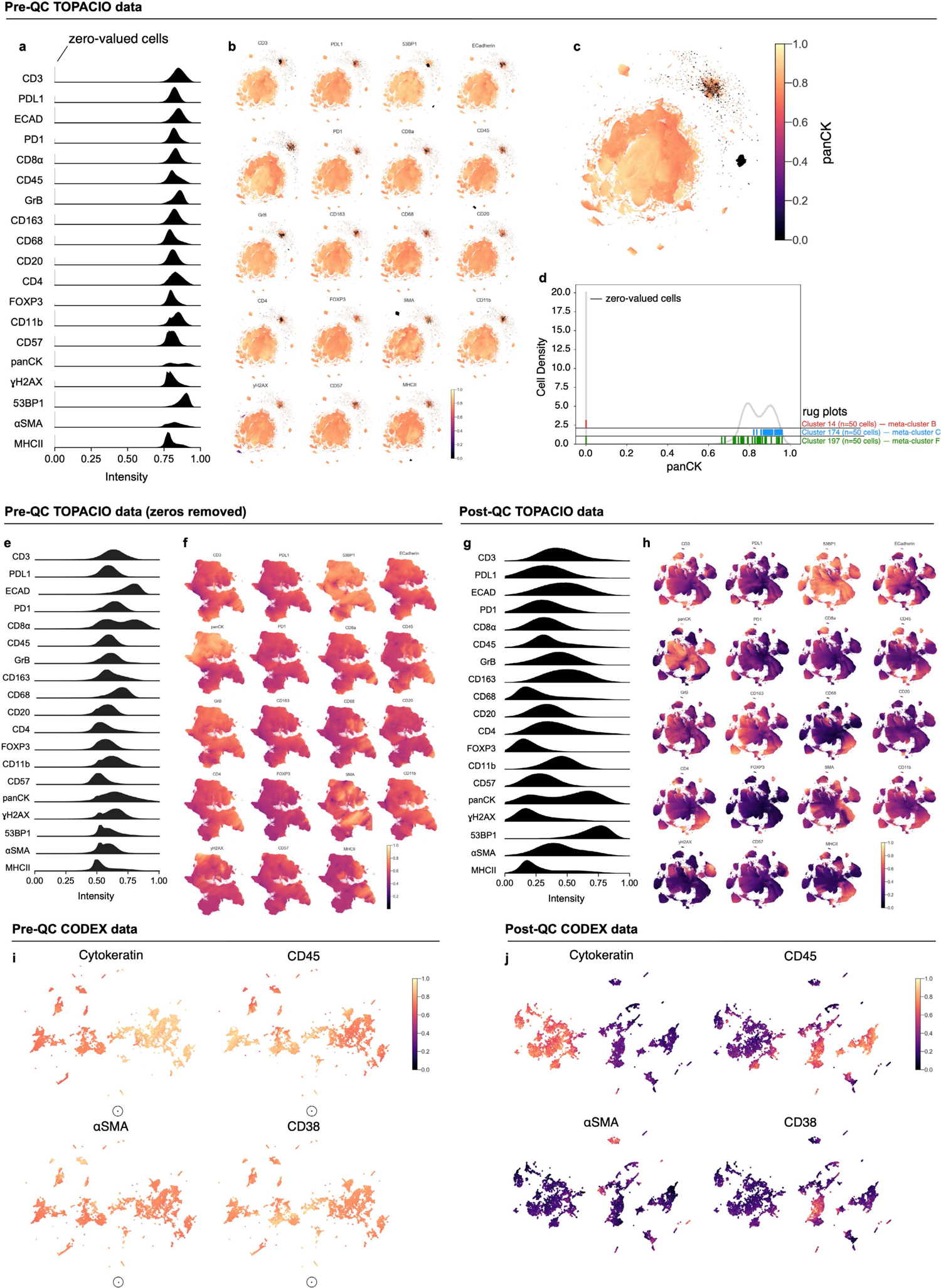
Impact of image background subtraction on derived single-cell data. **a**, Ridge plots showing the distribution of cells according to channel signal intensities in the pre-QC TOPACIO dataset showing the presence of zero-valued cells in each channel (vertical lines at far left). **b**, Channel colormaps applied to cells in the pre-QC TOPACIO embedding showing the presence of small, dark clusters corresponding to cells with at or near-zero signal intensities in the corresponding channel which by contrast makes all other cells appear bright for a given marker. **c**, PanCK channel from panel (b) enlarged to show detail. **d**, Histogram distribution of cells in the pre-QC TOPACIO dataset according to panCK signal. Rugplot plots (vertical ticks at bottom of histogram) show where randomly selected cells from cluster 14 (meta-cluster B, red), cluster 174 (meta-cluster C, blue), and cluster 197 (meta-cluster F, green) reside in the distribution. **e**, Ridge plots showing the distribution of cells according to channel signal intensities in the pre-QC TOPACIO dataset after the removal of zero-valued cells. **f,** Channel colormaps applied to cells in the pre-QC TOPACIO embedding after the removal of zero-valued cells showing that small, dark populations of cells are abrogated by the removal of zero-valued outliers. **g**, Ridge plots showing the distribution of cells according to channel signal intensities in the post-QC TOPACIO dataset allowing the natural distribution of signals to become apparent. **h,** Channel colormaps applied to cells in the post-QC TOPACIO embedding showing high contrast between populations of immunonegative and immunopositive cells for each marker. **i**, Channel colormaps applied to cells in the pre-QC CODEX embedding (Dataset 6) showing scant dim outliers (circles) which, in contrast, make other cells in the embedding appear bright for each marker. See **Online Supplementary** Fig. 9 for full set of marker channels. **j**, Channel colormaps applied to cells in the post-QC CODEX embedding showing high contrast between immunopositive and immunonegative cell populations cells after dim outliers have been removed. See **Online Supplementary** Fig. 10 for full set of marker channels.

**Supplementary Fig. 3.**
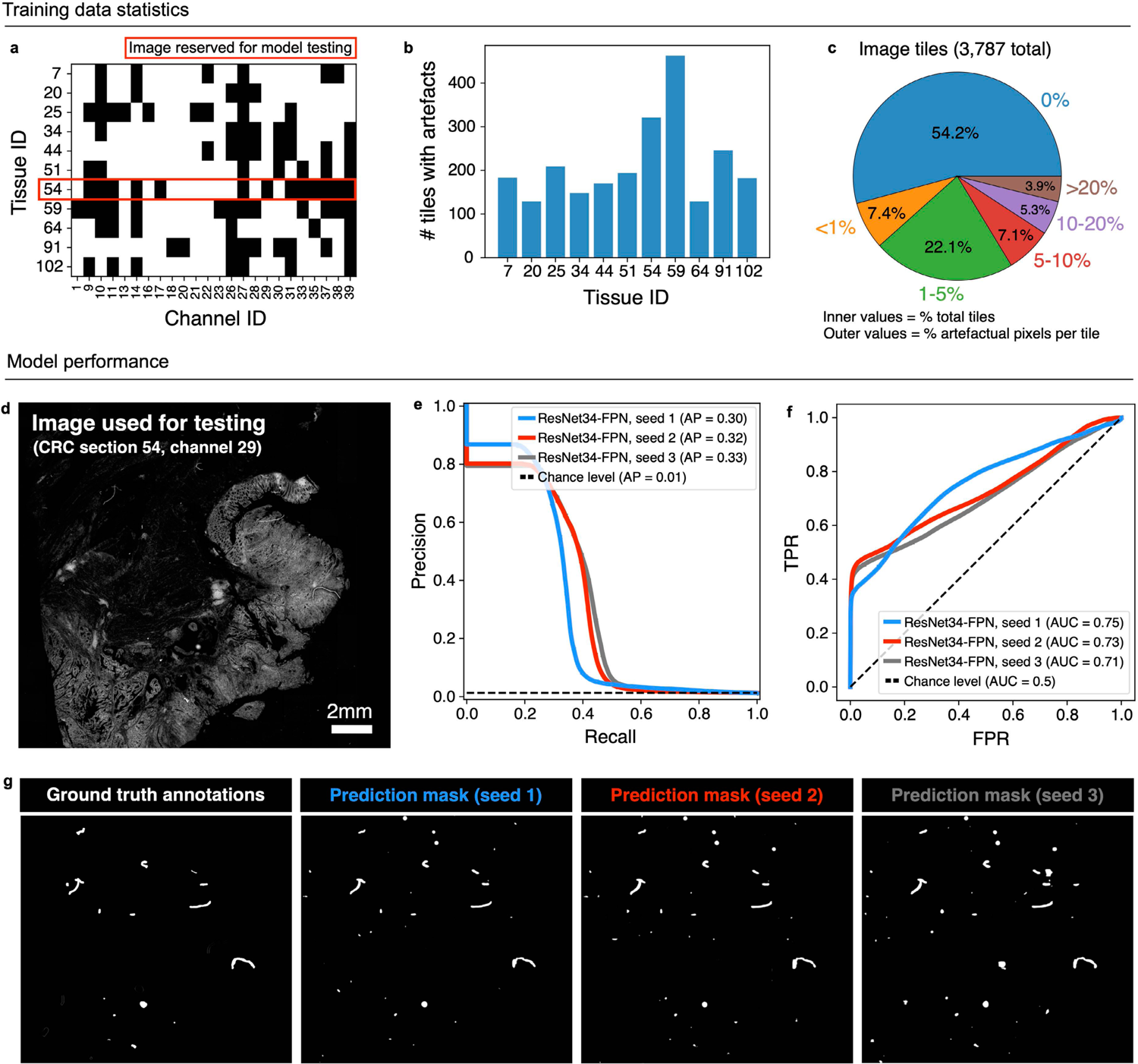
Developing a DL model for automated artefact detection tissue. **a**, Binary matrix showing the channels impacted by visual artefacts (e.g., illumination aberrations, slide debris, etc.) in 11 sections of the same CRC specimen. **b**, Barchart showing the number of 2048×2048-pixel image tiles affected by artefacts per tissue section. **c**, Pie chart showing the percentage of image tiles used for model training and validation (inner percentages) containing different percentages of artefactual pixels (outer percentages). **d**, CRC tissue section 54, channel 29 was used for model testing. **e**, Precision-recall plot showing the average precision (AP) for three replicates of the ResNet-FPN model architecture whose FPN network was initialized with different model weights to evaluate technical reproducibility. **f**, Receiver operating characteristic (ROC) curve showing the area under the curve (AUC) values for the same three replicates of the ResNet-FPN model shown in panel (e). **g**, Ground truth artefact mask (far left) and predicted artefact masks from the three replicate ResNet-FPN models.

## Notes

https://labsyspharm.github.io/cylinter

https://github.com/labsyspharm/cylinter

https://github.com/labsyspharm/cylinter-paper

https://www.synapse.org/#!Synapse:syn24193163/wiki/624232

### Summary of Updates

This version reflects substantial textual and figure edits prompted by peer-review. These include additional data analysis, a description of a deep-learning model for automated artifact detection, and multiple associated modifications to the CyLinter codebase, project website, and supporting Wiki page. We believe that these revisions substantially improve the content, readability, and potential impact of our manuscript.

